# Personalized mapping of inhibitory spinal cord circuits *in vivo* via non-invasive neural decoding and *in silico* modelling

**DOI:** 10.1101/2025.03.25.645235

**Authors:** Alejandro Pascual-Valdunciel, Natalia T. Cónsul, Robert M. Brownstone, Marco Beato, Dario Farina, Filipe Nascimento, M. Görkem Özyurt

## Abstract

Studying human motoneuron activity through electromyography (EMG) can yield insights into the operation of fundamental spinal cord microcircuits. Traditional surface and needle electromyography (EMG) methodologies have limited capacity to shed light on the diversity of motor unit (MU) control strategies that may be unique to each individual. Here, we employed high-density surface EMG (HDsEMG) to sample multiple MUs per subject to investigate the dynamics of inhibitory spinal microcircuits in both upper and lower limb control. We characterised the net inhibition as a function of individual MU firing rates, revealing subject-specific relationships. *In silico* modelling replicated these experimental characteristics and suggested that properties of the inhibitory currents rather than motoneuron size are responsible for net functional inhibition. Our results show that HDsEMG can highlight distinct control strategies across circuits and motor pools, revealing subject-specific properties of inhibitory spinal microcircuits.

**TEASER:** High-density surface EMG electrodes can reveal the functional properties of inhibitory spinal circuits

## INTRODUCTION

Movement relies on the precise coordination of muscle contractions, enabling fundamental actions like grasping and walking. Muscles are composed of individual fibres innervated by spinal motoneurons. A single motoneuron connects to a group of muscle fibres, collectively forming a motor unit (MU). While large parts of the central nervous system are responsible for various aspects of movement, motoneurons serve as the “final common pathway” responsible for its execution^1^.

Electromyography (EMG) is a valuable tool for studying motoneurons and their associated circuits. EMG can be used to record gross muscle activity, reflecting neuronal activity in the population (pool) of motoneurons innervating that muscle, through global surface (sEMG) or intramuscular EMG (iEMG). Additionally, the activity of individual MUs can be recorded via single unit iEMG, which reflects the action potentials of individual motoneurons. As such, motoneurons are the most readily recordable cell type in the human central nervous system (CNS), offering a valuable window into CNS activity.

The region of the CNS that is responsible for organising the pattern of muscle contractions for limb movement is the spinal cord. Various fundamental spinal circuits have been identified in animal studies, with some also characterised in humans. One well-characterised example is the circuit controlling reciprocal inhibition between antagonist muscles, mediated by Ia inhibitory interneurons^2,3^. This circuit, also responsible for inhibition of antagonist muscles in stretch reflexes, plays a key role in ensuring proper flexor-extensor alternation during locomotion^4^. Another example is the circuit responsible for producing a “cutaneous silent period” (CSP)^5^, a transient pause in voluntary muscle contraction produced by electrical stimulation of a cutaneous nerve, part of a protective behaviour in response to noxious cutaneous stimuli.

While both of these circuits can be measured in humans, current techniques fail to give an accurate picture of their operation. In humans, sEMG recordings provide low-resolution data and underestimate the duration of both circuits^6–8^. On the other hand, iEMG using fine wires or needles provides single unit resolution and a better estimation of inhibition duration but is invasive and only samples one or few MUs within highly heterogeneous motoneuron populations, and therefore falls short in estimating the temporal characteristics of circuits as a whole. To overcome the limited sampling, subject data are usually pooled^9^. But doing so relies on the assumption that motor pool dynamics are similar between individuals^10^, which we know is not the case^10,11^. That is, only limited mechanistic, physiological insights into the operation of spinal circuits can be gained using sEMG and iEMG methods.

If, as in animal studies^12^, the function of human inhibitory circuits reflects the status of a disease such as amyotrophic lateral sclerosis (ALS), then it will be necessary to measure inhibition accurately, reproducibly, and in a subject-specific manner. Indeed, recent studies have pointed to spinal circuit alterations in neurological diseases^13–15^, yet the technical constraints imposed by traditional sEMG and iEMG have meant the loss of subject-specificity. Enhanced sampling methods capable of sampling multiple MUs and capturing their respective spinal cord circuit dynamics will greatly improve the accuracy of physiological and pathophysiological measures. We have therefore sought to establish new methodologies for probing spinal circuit function at higher resolution, which would enable more precise assessments of MU behaviour and individual variability.

Advancements in high-density surface EMG (HDsEMG) techniques are transforming the landscape in the field of motor physiology^16^. This non-invasive method uses arrays of closely-spaced electrodes to achieve high spatial and temporal resolution of the activity of individual MUs, recording a high proportion of units that comprise an individual muscle. HDsEMG has facilitated the study of a variety of MU properties, such as conduction velocity and discharge rate^17^, as well as network-level features such as common synaptic inputs to motoneurons during movements, indicating a potential to shed light into the mechanisms of motoneuron integration of input and recruitment across a pool^18,19^.

In this work, we present an optimized framework for the use of HDsEMG in the study of spinal circuits in both upper and lower limb muscles. We focus on two distinct inhibitory pathways, CSP and reciprocal inhibition, as these circuits are involved in key physiological functions and have biomarker potential^13,20,21^. Using simultaneous testing across different muscles, we draw inferences about distinct inhibitory spinal pathways. By successfully sampling multiple MUs per subject, we were able to conduct subject-specific analyses of the temporal dynamics of CSP and reciprocal inhibition. This allowed us to characterise these circuits by taking subject-level variations into account and to provide more accurate estimates of functional inhibition acting on firing motoneurons. We then modelled these circuits *in silico* to obtain insights into the relationship between synaptic inhibition and motoneuron intrinsic properties. Together, we provide a blueprint for the use of HDsEMG to detail spinal cord circuit mechanisms and to further validate its application in neuromuscular research and clinics.

## RESULTS

### Subject-specific analysis of Cutaneous Silent Period (CSP)

Reasoning that the use of HDsEMG provides an opportunity to study stimulus-evoked inhibition of many motoneurons simultaneously, and that the duration of inhibition reflects the temporal profile of the net inhibitory postsynaptic potential received by each motoneuron^7,22^, we turned to methods of quantifying the times of onset and termination of inhibition based on spiking profiles. Peristimulus time histograms (PSTHs) constructed from single unit data show spiking activity in response to the stimulus and accurately reflect the onset latency of inhibition of a MU^23,24^. Plotting the frequency of firing, peristimulus frequencygrams (PSFs) can pinpoint when the firing frequency returns to its baseline (prestimulus) level, reflecting the termination of inhibition of firing^7,13,22^. Thus, from these two plots, the duration of “functional inhibition” can be calculated^7,22,25^. We then used PSTH and PSF measures for all units recorded via HDsEMG, including all MUs with significant inhibition defined as a deviation of the CUSUM below the limits seen in the 200ms preceding the stimulus (<u>Fig. S1</u>).

We initially focused on the upper limb CSP, a robust and reproducible stimulus carried by A-delta fibres to spinal dorsal interneurons, which in turn inhibit motoneurons innervating intrinsic hand muscles^5,26–28^ (<u>Fig 1A</u>). We placed 64-channel HDsEMG grids on top of the first dorsal interosseous (FDI) muscle and instructed the participants to perform a pinching task for 200s at 10% of their maximum voluntary contraction (MVC), electrically stimulating the fifth finger (at x10 the sensory threshold) every 1.8s^27^. Single MUs were decomposed from the raw EMG (<u>Fig 1A</u>) to measure the temporal properties of inhibition (<u>Fig. 1B</u>).

**Fig. 1.**
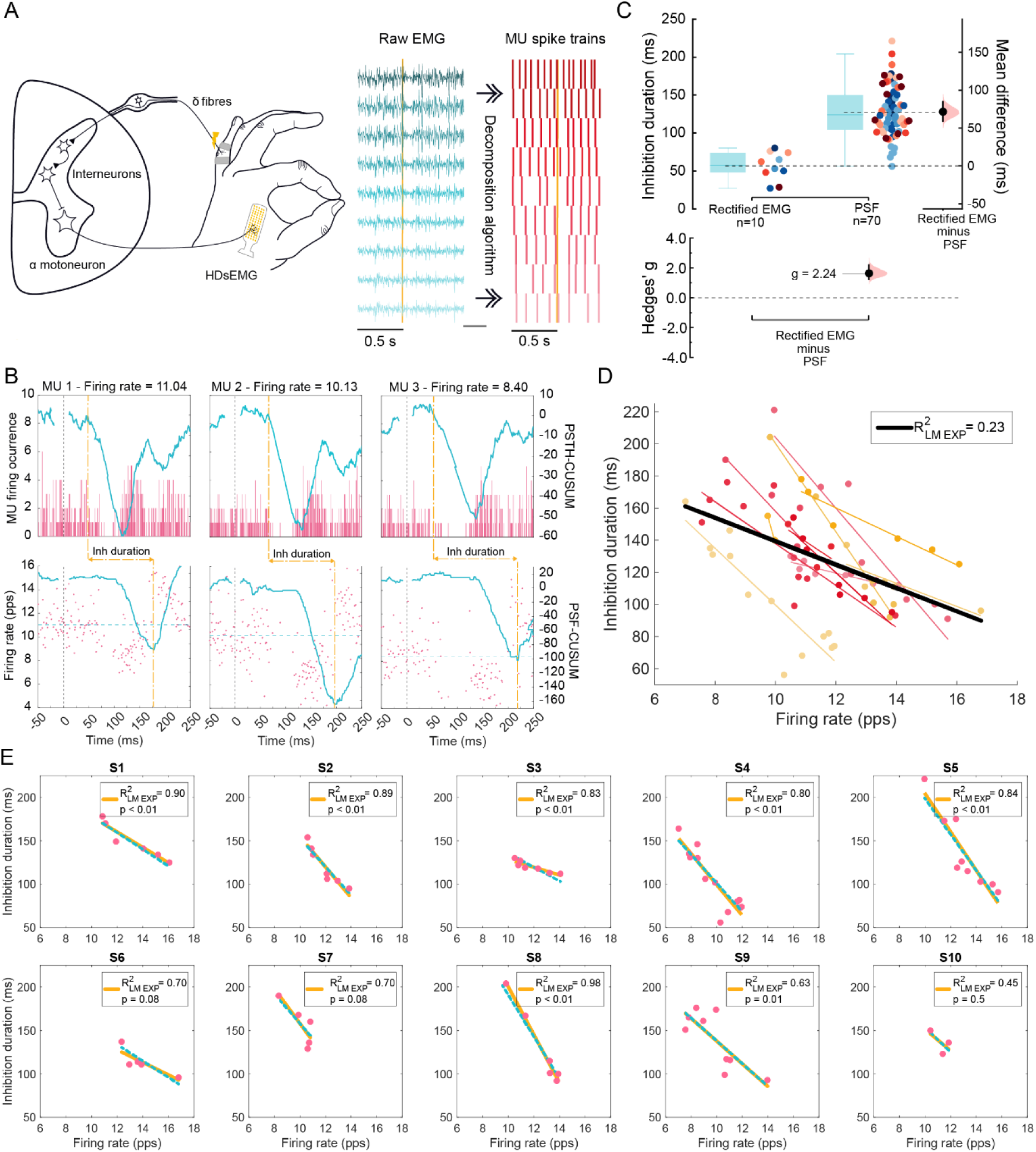
Characterization of CSP on individual motoneurons in humans. **(A)** Schematic of the CSP spinal pathway and experimental setup (left) and the decomposition methods used to extract the firing phase and frequency of individual MUs with HDsEMG (right). **(B)** Examples of PSTH (top) and PSF (bottom) of individual MUs from the FDI muscle active at different frequencies, with cumulative summation (CUSUM) traces in blue for each. Dotted yellow lines depicted inhibition start defined by PSTH and end by PSF. **(C)** Estimation plots illustrating the comparison of the inhibition duration estimation methods: rectified sEMG and individual MUs with the PSF method, with dots colour coded per individual. **(D)** Estimation of inhibition duration of single MUs and their firing rate across subjects. Coloured dots represent MUs and the linear regression fit per individual; black line represents the linear regression fit for the group data with respective R^2^ shown. **(E)** Inhibition duration of single MUs vs their firing rate per individual (pink dots). Yellow lines represent the individualized linear regression models (LM_exp_) with respective R^2^ and p-value; blue dotted lines represent the linear mixed model (LMM) regression per subject; pps – pulse per second.

In addition to the decomposition of individual MUs, the HDsEMG grid can also act as a standard bipolar sEMG electrode by recording the voltage difference between two adjacent electrodes, providing a conventional sEMG signal for assessing overall muscle activity (<u>Fig. S1</u>). We therefore compared the duration of inhibition obtained from the individual MUs decomposed with HDsEMG to the conventional sEMG signal obtained during the same recordings (<u>Fig 1C</u>). We found a consistently shorter CSP duration when measured from the rectified sEMG (58±18ms) compared to the durations measured from individual MUs using PSF (121±22ms), indicating that sEMG recordings are sensitive only to the shortest duration inhibitory responses (µ_diff_ = 72ms, 95% CI = [59,85]; *g* = 2.25, 95% CI = [1.82, 2.84]). The sEMG provides an averaged signal of multiple MUs, each with slightly different timing for inhibition onset and termination. Because sEMG estimation of inhibition is defined solely by spike occurrence or spike count^7^, MUs with shorter inhibition periods resume firing earlier, potentially masking ongoing inhibition in other units that experience longer duration inhibition. Moreover, even when a MU resumes firing, its discharge rate may remain below baseline reflecting a continued hyperpolarization of the membrane potential. As a result, PSF estimates better reflect the underlying membrane dynamics and thus capture longer inhibition durations by incorporating both the presence and amplitude of the inhibitory postsynaptic potentials^29^. Overall, single unit HDsEMG data revealed a wide range of durations, with many units being inhibited over twice as long as estimated by rectified global sEMG.

The duration of the CSP during voluntary contractions is highly variable and influenced by the MU discharge rate^6,20^. Therefore, interpreting inhibition duration is best done relative to firing rate^6,20^. The coefficient of determination (R^2^) is a standard metric for assessing goodness-of-fit^30,31^, and is widely used to validate linear relationships, such as firing rate *versus* inhibition duration in motoneuron studies^7,13,14,20^. A R² value of 1 indicates a perfect linear relationship, meaning that firing rate fully accounts for the variability in inhibition duration, whereas an R² of 0 means that firing rate provides no explanatory power, with inhibition duration varying independently. For this study we classified the strength of the R^2^ using thresholds of 0.67, 0.33, or 0.19, corresponding to strong, moderate and weak relationships, respectively, based on established statistical conventions^32,33^ (<u><u>Fig. S2</u></u>). Additionally, we report p-values to assess whether the relationship between firing rate and inhibition duration is statistically significant.

When plotting firing rate against CSP duration, we observed that inhibition duration is poorly predicted by firing rate for all MUs combined from all participants, as shown by a weak R^2^ (<u>Fig. 1D</u>). However, this R^2^ (R^2^=0.24) was similar to that reported previously using iEMG for healthy subjects (R^2^=0.31)^13^. Although low R^2^ values may still help identify trends within the data, they are limited in predictive accuracy and precision^34^.

While single unit iEMG has been useful to establish group-level clinical outcomes when pooling data from groups of subjects^13,14^, at best it can only sample a few MUs from a single subject. This limited sampling of the motor pool within a single individual fails to capture motoneuron heterogeneity within and between subjects, resulting in low-dimensional data that reduce measurement precision and limit the ability to make subject-specific assessments^9^. In contrast, the increased MU sampling enabled by HDsEMG allows for the creation of robust hierarchical datasets (level 1 - MUs, level 2 - subjects), making it possible to assess how individual differences influence the relationship between inhibition duration and discharge rate for CSP.

To quantify the influence of subject-specific variability in the interpretation of CSP experimental data, we implemented a linear model (LM_exp_) for each subject and employed a linear mixed model (LMM) to account for subject-specific random effects. For the FDI muscle, individual LM_exp_ consistently outperformed the pooled correlation for firing rate *vs* inhibition duration relationships, yielding higher R^2^ values. Specifically, 6 of the 10 subjects exhibited strong R^2^, indicating that firing rate explained a substantial portion of the variability in inhibition duration (p<0.05). An additional 2 subjects had a high R^2^ but p=0.08, potentially reflecting low power due to the few MUs extracted. The remaining 2 individuals exhibited moderate R^2^ with the subject with the fewest sampled MUs (3 MUs) showing no meaningful relationship (p=0.5) (<u>Fig. 1E</u>). The LMM results indicate that inhibition duration varied significantly with firing rate, with a decrease of 15ms in inhibition duration for every 1Hz increase in discharge rate (<u>table S1</u>). But more importantly, the LMM captured 84% of data variability (R² =0.84), driven by inter-subject differences that accounted for 80% of total variance (ICC=0.80). The LMM analysis revealed significant differences in inhibitory profiles across subjects, with subjects with longer inhibition durations having less reduction in inhibition duration with increasing firing rate (see <u>table S1</u>). The disparities in the measured effects between grouped *versus* subject-specific analyses, underscore the importance of considering individuality when interpreting the effect of synaptic inhibition on MU behaviour.

Taken together, we demonstrate that HDsEMG is a more sensitive method than traditional EMG approaches for estimating CSP duration. Unlike traditional methods, HDsEMG offers a non-invasive estimation comparable to single unit iEMG, while also allowing for the sampling of multiple MUs per subject. Furthermore, the hierarchical datasets generated through HDsEMG provide a robust framework for studying inhibition duration on a subject-specific level, accounting for inter-individual variability.

### Predicting CSP characteristics using *in silico* modelling

Inhibition of motoneurons depends on the strength of the inhibitory inputs and the response properties of the motoneurons. Intrinsic neuronal properties influence how synaptic inputs are integrated, potentially impacting inhibition duration. For example, the physical dimensions of motoneurons are directly linked to their intrinsic properties: motoneurons with larger somas and dendritic trees require greater synaptic input to achieve the same membrane voltage change as smaller motoneurons^35,36^.

Historically, *in vivo* and *in vitro* preparations from animals have enabled single-cell recordings, allowing for controlled current injections and intracellular voltage measurements^37–40^. These techniques provide insight into synaptic inputs and intrinsic properties, such as whole cell capacitance, which can be used as a surrogate marker of motoneuron size. HDsEMG, however, cannot directly measure these parameters.

During voluntary contractions, FDI motoneurons firing at higher firing rates exhibit shorter CSP inhibition, despite delivering the same nerve stimulation intensity, indicating that motoneurons likely receive the same synaptic drive from afferents^35,36,41^. Is this effect purely due to firing rate, and/or do differences in intrinsic properties play a role? To disentangle these factors, we built an *in silico* model replicating FDI CSP inhibition, and manipulated motoneuron size and firing rate to assess their distinct effects on inhibition duration.

We generated a two-compartment Hodgkin-Huxley biophysical model^42^ and simulated the CSP experiments previously described. The model simulated common input to a motor pool of 20 motoneurons replicating an isometric contraction, with an inhibitory synaptic input delivered every 1.8s during the steady phase of the isometric contraction (<u>Fig. 2A</u>). The motoneurons were modelled based on the characteristics of S-type motoneurons, which primarily govern sustained, low-force contractions (<u>Table S6</u>). These motoneurons were assigned sizes ranging from the smallest to the largest (189-214pF of cell soma capacitance), with properties linearly interpolated to reflect a physiological gradient in excitability and recruitment thresholds, consistent with known motor pool organization^35,43,44^. We recorded the spike train outputs in all motoneurons across 154 realizations, each defined by a unique combination of inhibitory amplitude (A; 1-3 arbitrary units [a.u.], in 0.2a.u. steps) and duration (τ; 7– 20ms, in 1ms steps), distributed uniformly across motoneurons. For each realization, we quantified inhibition duration of each simulated motoneuron using the same PSTH and PSF methods applied to the experimental HDsEMG data (<u>Fig. 2B</u>), and kept only the units with significant inhibition based on the CUSUM approach described above for HDsEMG recordings (<u>Fig. S1</u>). The simulated inhibition durations and respective firing rates were linearly fitted for each realization, leading to 154 linear models (LM_sim_) with a range of R^2^, slopes and intercept consistent with those observed across subjects (<u>Fig. S3A</u>).

**Fig. 2.**
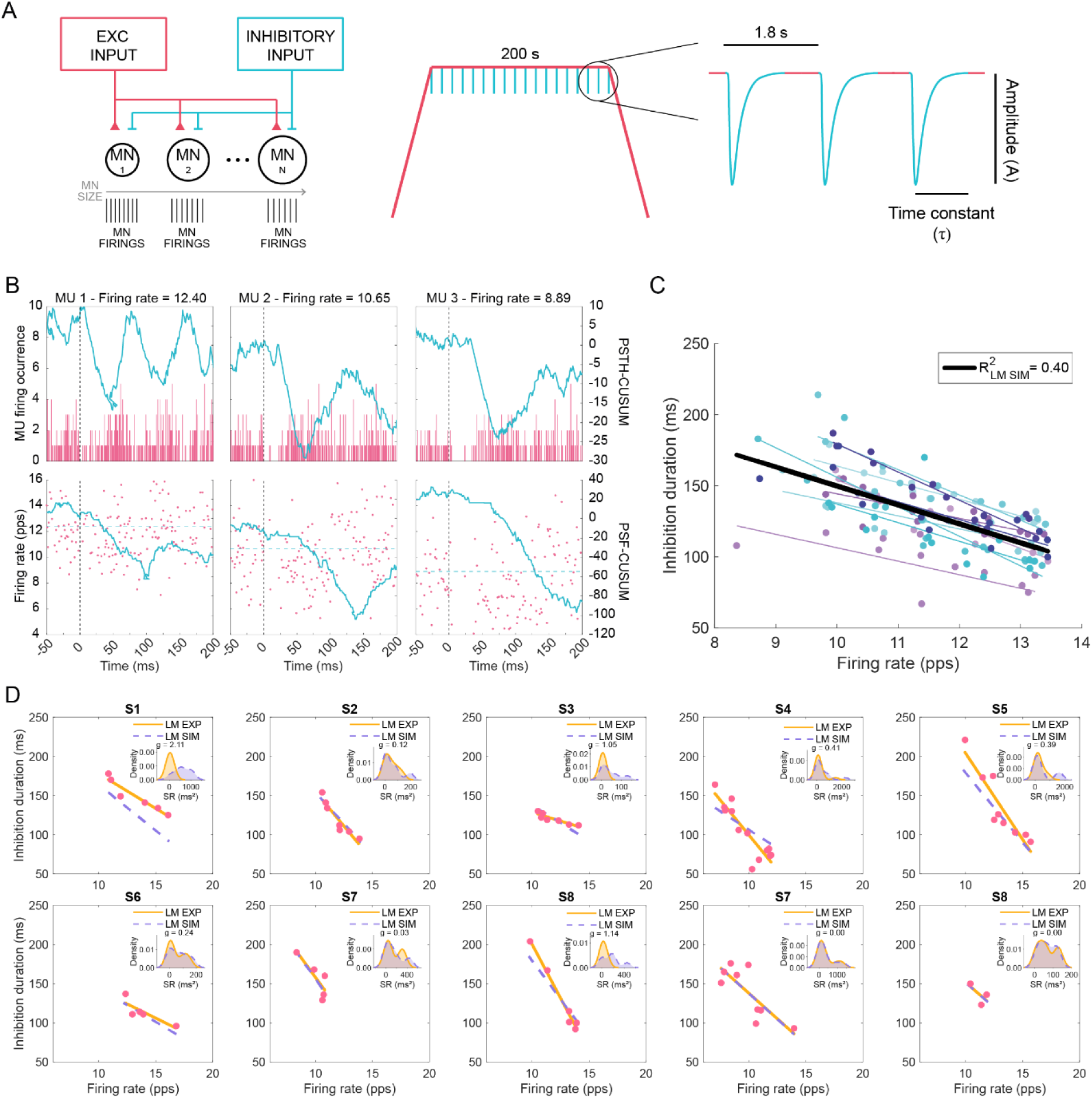
Characterization of CSP on individual motoneurons with computational modelling. **(A)** Schematic of the *in silico* computational model depicting the common excitatory input and inhibitory input delivered to motoneurons (MN) of varying size (left), used to simulate the experimental MU firing during voluntary contraction with periodic inhibitory drive to motoneurons delivered during the plateau phase (right). **(B)** Examples of PSTH (top) and PSF (bottom) of simulated MUs at different frequencies, with cumulative summation (CUSUM) traces in blue. **(C)** Estimation of inhibition duration of single MUs and their firing rate across the LM_sim_ optimized for each subject. Individual simulated MUs and each of the LM_sim_ fits are shown in colour; black line represents the linear regression fit for the simulated group data with respective R^2^ shown. **(D)** Experimental MU firing rate and inhibition duration (pink dots) with respective regression line (orange line; same LM_exp_ as in <u>Fig. 1E</u>) and optimal LM_sim_ fit for each subject (dotted purple line). Inset plots show distributions of square of residuals (SR) as kernel density estimates along with the corresponding Hedges’ *g* value; pps – pulse per second.

We then identified the LM_sim_ best suited to replicate the experimentally observed dependency of inhibition duration on firing rate for each of the 10 subjects. We inputted each subject’s experimental firing rates into all 154 LM_sim_ formulas, and selected the LM_sim_ with the lowest mean squared error (MSE), which minimized errors between predicted and observed inhibition durations. The LM_sim_ with the lowest MSE was selected as the best-fitting model for that subject, ensuring minimal prediction error at the individual level (Figs. <u>S2D</u> and <u>S3B</u>; <u>table S2</u>).

To illustrate comparisons between experimental data fits (LM_exp_) and the LM_sim_ optimized for each of the 10 subjects (<u>Fig. 2C</u>), we analysed the distributions of square of residuals between the fits relative to each of the experimental datapoints and calculated Hedges *g*’ to assess differences between the distributions (<u>Fig. 2D</u> and <u>S3C</u>). Apart from subjects 1, 3 and 8 who showed a notable discrepancy between the residuals, effect sizes for the remaining subjects were small (0.15<g<0.50) or negligible (g<0.15). These results suggest that, while the computational model generally captured observed CSP inhibition duration dependency on firing rate, its performance varied across individuals.

### Motoneuron size does not contribute to variations in inhibition duration according to biophysical modelling

Motoneurons vary in size and form a diverse group, even within the same motor pool. This structural variation leads to a graded recruitment pattern in response to common input, where smaller, low threshold motoneurons reach their action potential threshold before larger ones. As the input increases, smaller motoneurons will fire faster at the time that the larger motoneurons are recruited (and fire at low rates). Our experimental and simulated results indicate that the duration of functional inhibition is longer in later recruited motoneurons, which fire slower (towards the left-end of the regression panels). Yet, it is unclear why the inhibition is longer in these neurons: is it because they are larger in size (late recruited motoneurons are usually larger motoneurons) or because they fire at a lower rate (i.e. receiving lower net excitatory input which may cancel out inhibition)? We cannot answer this question experimentally, so we used our *in silico* model to investigate whether motoneuron size influences variations in inhibition duration.

We simulated motoneurons using the optimized model parameters from subject 8, characterized by long-duration (19ms) and high-amplitude (2.8 a.u.) simulated inhibitory inputs that produced a strong dependency of inhibition with firing rate (table S2). We also ran a secondary set of simulations using parameters from subject 4, which differed substantially in both the optimal parameter values for synaptic input amplitude (1.8a.u.) and duration (7ms) and the linear fit (table S2). These contrasting conditions provided an opportunity to explore the effects of motoneuron size under distinctly different simulation conditions. For these simulations, we doubled the simulated motoneuron pool (40 instead of 20) to improve resolution in characterizing intrinsic property gradients. The common input was calibrated to generate steady firing rates of 11-14Hz during the plateau phase of the ramp (0.2 to 0.28a.u. in 0.01a.u. steps). Synaptic inputs, fixed in amplitude (1.8a.u. or 2.8a.u.) and duration (7ms or 19ms) to reflect physiological sensory spinal circuit behaviours^12,41,45^, were delivered every 1.8s, while motoneuron size was systematically varied based on S-type motoneuron parameters (table <u>S6</u>), as previously described the CSP (<u>Fig. 3A</u>). This approached enabled precise control of MU firing rates and manipulation of motoneuron dimensions within set ranges, isolating the effect of cell size (i.e. capacitance) on inhibition duration.

**Fig. 3.**
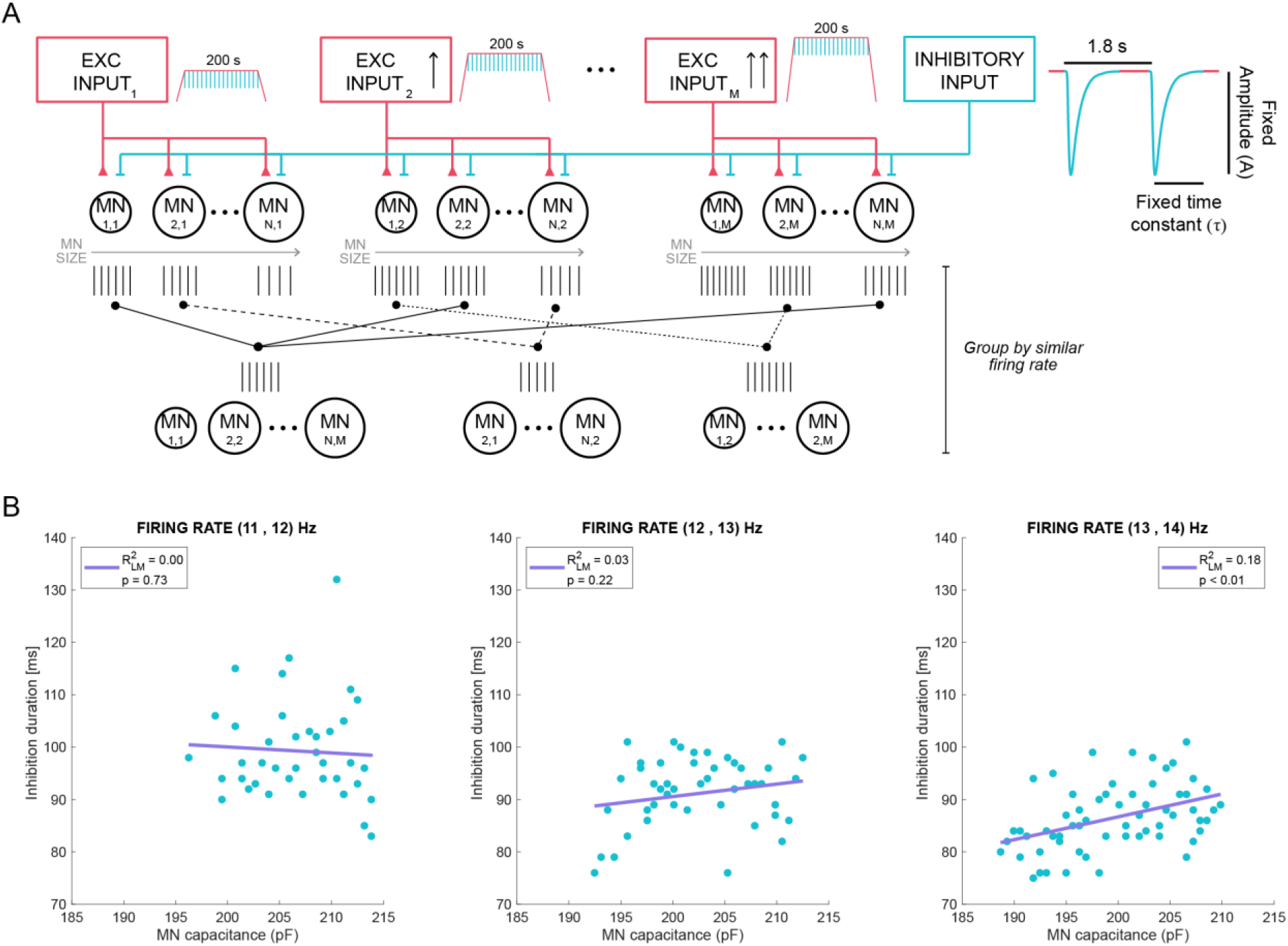
Influence of motoneuron size on inhibition duration *in silico*. **(A)** Schematic of the computational model, showing the different common inputs that generated 3 distinct firing rates, with motoneuron (MN) size varying per discharge rate group and with inhibition strength of fixed amplitude and duration. **(B)** Inhibition duration across simulated motoneurons with varying size (blue dots) with each plot representing a different firing rate, with purple line representing the linear regression between capacitance and inhibition duration. Capacitance values represent simulated soma capacitance. Simulations were obtained using optimized parameters for subject 8 (inhibitory input amplitude of 2.8a.u. and duration of 19ms).

To examine the effect of motoneuron size across different discharge rates, motoneurons were clustered into three different groups by firing rate (11-12Hz, 13-14Hz, 14-15Hz) and compared across soma sizes (<u>Fig. 3B</u> and <u>Fig. S4</u>). Group comparisons revealed that inhibition duration decreases as the firing rate increases (<u>table S3</u>). However, within each firing rate group, motoneuron capacitance did not explain variations in inhibition duration as shown by the very weak R^2^ values. These findings were similar for the two sets of optimizing parameters tested for A and τ (<u>Fig. 3B</u> and <u>Fig. S4</u>). And in fact, motoneuron capacitance did not introduce any variability to the relationship between inhibition duration and firing rate as shown by the negligible ICC (ICC_capacitance_ <0.20; <u>table S3</u>). These results suggest that, within the tightly controlled parameters of our *in silico* approach, modelled inhibition duration is predominantly driven by firing rate rather than motoneuron size, implying that active properties (e.g. firing rate) outweigh passive properties (e.g. capacitance) in shaping the effectiveness and duration of functional inhibition.

### Estimating reciprocal inhibition in the Tibialis Anterior with HDsEMG

To examine whether our finding that inhibition depends mainly on firing rate can be extrapolated to other inhibitory circuits, we next turned to the lower limb, with a focus on a different inhibitory circuit - reciprocal inhibition. This circuit involves recruitment of a different set (compared to CSP) of sensory afferents (group I) and spinal interneurons (Ia interneurons) and is responsible for flexor-extensor alternations during movement^8,46^. To study reciprocal inhibition, we tested 8 subjects by placing a 256 multi-electrode HDsEMG grid over the TA muscle, and evoked reciprocal inhibition from triceps surae afferent to TA motoneurons through tibial nerve stimulation (<u>Fig. 4A</u>). The temporal properties of inhibition were estimated as above using PSF and PSTH (<u>Fig. 4B</u>).

**Fig. 4.**
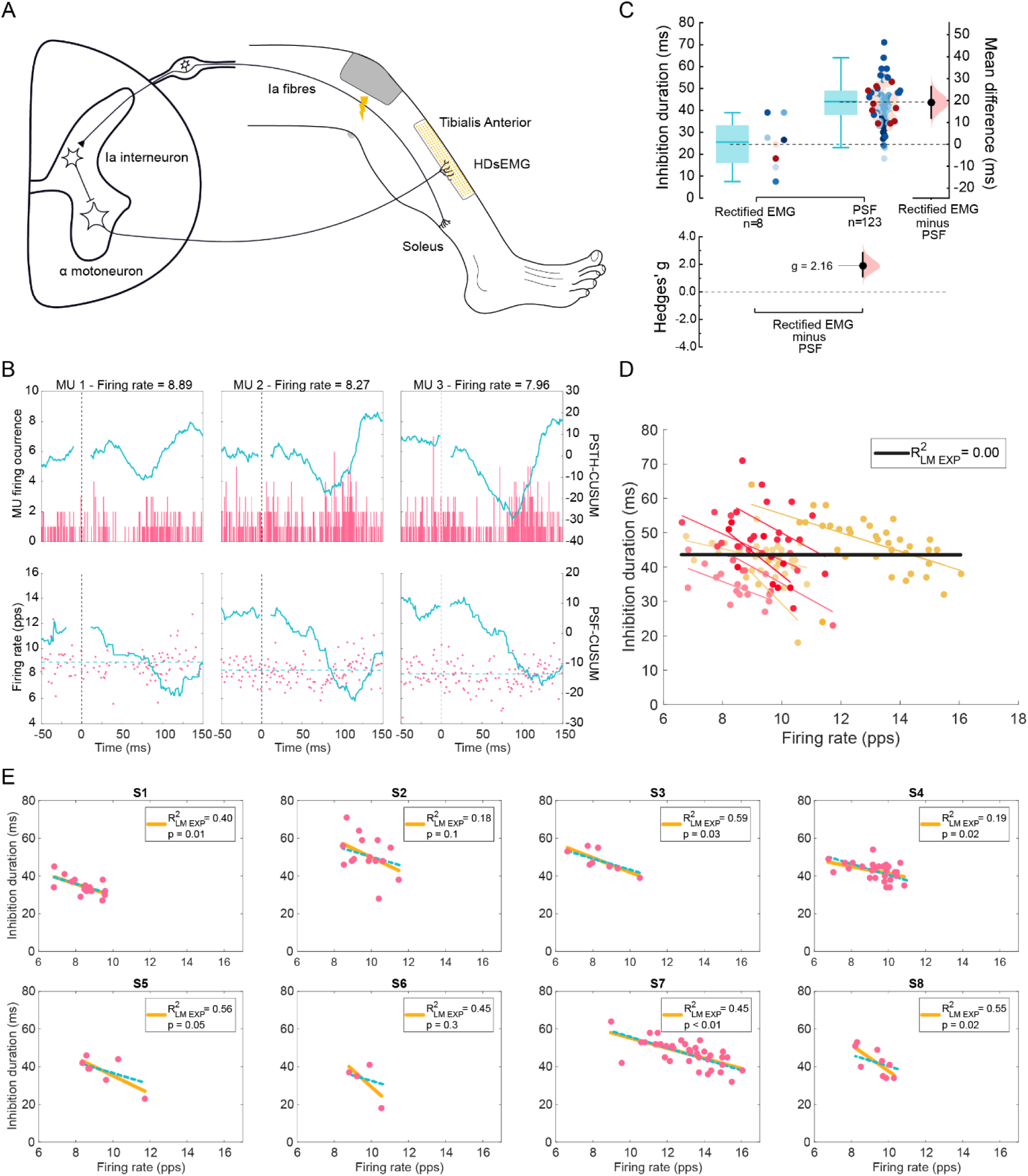
Characterization of reciprocal inhibition of the TA on individual motoneurons in humans. **(A)** Schematic of the reciprocal inhibition spinal circuit and experimental setup used to measure inhibition from the TA muscle. **(B)** Examples of PSTH (top) and PSF (bottom) of individual MUs at different frequencies following TA muscle activation, with cumulative summation (CUSUM) traces in blue for each. **(C)** Estimation plots depicting comparison of the inhibition duration estimation methods: rectified surface EMG and individual MUs with the PSTH-PSF method. **(D)** Estimation of inhibition duration of single MUs with varying firing rate across subjects. Coloured dots represent MUs and the linear regression fit per individual; black line represents the linear regression fit for the group data with respective R^2^ shown. **(E)** Inhibition duration of single MUs and their firing rate per individual (pink dots). Yellow lines represent the individualized linear fits (LM_exp_) with respective R^2^ and p-value; blue dotted lines represent the LMM regression per subject; pps – pulse per second.

We first compared the estimates of inhibition duration between the individual MUs decomposed through HDsEMG and the conventional sEMG signal obtained during the same recordings. Similar to our findings for CSP, sEMG measurements underestimated the duration of reciprocal inhibition (25±11ms), compared to the inhibition of individual MUs (42±7ms) obtained using HDsEMG (µ_diff_ = 19ms, 95% CI = [12,26]; *g* = 2.16, 95% CI = [1.35, 3.13]); <u>Fig 4C</u>). This again highlights that individual MUs sampled with HDsEMG provide a wider range of inhibition durations.

Given that inhibition duration varies with discharge rate for CSP, we examined the dependence of reciprocal inhibition on motoneuron firing rates. Pooled MU data across subjects showed that inhibition duration is poorly predicted by firing rate when aggregating MU data from multiple subjects (R^2^=0.00, <u>Fig. 4D</u>). On the other hand, individual LM_exp_ yielded a moderate R^2^ for 4 of the 8 subjects (1, 3, 4 and 8 p<0.05). For another subject, borderline-weak R^2^ was observed (R^2^=0.19, p=0.02). Subject 5 had a moderate R^2^ but a p=0.05, whereas subjects 2 and 6 did not have a statistically significant relationship. A LMM with random intercept revealed a decrease of 3ms in inhibition duration for every 1Hz increase in discharge rate, with inter-subject variability reflected by a R^2^ of 0.56 and an ICC_subject_ of 0.60 (<u>table S4</u>). These results highlight the importance of accounting for inter-subject variability to accurately interpret the relationship between discharge rate and duration of reciprocal inhibition.

Overall, individual LM_exp_ suggest a weaker relationship between reciprocal inhibition duration of TA MUs and discharge rate compared to that seen in CSP duration in FDI MUs. Although we sampled an average of 15 MUs for reciprocal inhibition and 7 MUs for CSP, firing rate tended to exhibit less explanatory power for reciprocal inhibition LM_exp_ (4 moderate and 1 weak R^2^; p<0.05) than for CSP LM_exp_ (6 strong and 1 moderate R^2^; p<0.05). For CSP, individuals with fewer sampled units generally exhibited lower R^2^ and higher p-values (e.g. subject 10, <u>Fig. 1D</u>). However, this pattern was not observed for reciprocal inhibition, where some participants with many sampled units still showed a weak R^2^ and, in some cases, no statistically significant linear relationship (e.g. subject 2, <u>Fig. 4D</u>).

Reciprocal inhibition measured from the TA had a ∼4-fold shorter inhibition duration and a 5-fold smaller change in inhibition duration with discharge rate than the CSP for FDI (tables <u>S1</u> and <u>S4</u>). This could impact the relationship between firing rate and inhibition duration by reducing the overall influence of inhibition on motoneuron discharge dynamics, potentially contributing to a weaker correlation between reciprocal inhibition duration and firing rate in TA MUs compared to CSP duration in FDI MUs. Nevertheless, similar to the experimental observations for the CSP, our results highlight the need to sample multiple units per subject to capture individual reciprocal inhibition effects.

### Predicting reciprocal inhibition characteristics using *in silico* modelling

We now extended our *in silico* approach to investigate reciprocal inhibition. We simulated the common input into a motor pool of 20 motoneurons of varying size (same as for CSP; table <u>S6</u>) to reproduce an isometric contraction. Initially, we ran simulations with a 0.20a.u. common input, which replicated experimental firing rates. However, to minimize background noise that affected PSTH-PSF inhibition duration estimates for some realizations, we performed additional simulations with a reduced 0.15a.u. common input. This adjustment preserved realistic firing rates while reducing noise interference when estimating inhibition duration. Inhibitory synaptic inputs received by motoneurons during the steady-state phase of the isometric contractions were delivered every 2s (<u>Fig. 5A</u>). Motoneuron size was systematically varied using the same parameters employed in earlier simulations (<u>table S6</u>). We recorded spike discharges across realizations, each defined by a combination of inhibitory inputs of varying A (1-3a.u. in 0.2 steps) and τ (1-4ms in 1ms steps). We analysed inhibition duration of each simulated motoneuron using the PSTH and PSF, keeping only units with detectable inhibition (<u>Fig. 5B</u>). The simulated firing rates and inhibition durations were fitted per realization (<u>Fig. S5</u>). For some realizations we obtained 2 or fewer units with detectable inhibition, which did not allow us to establish linear regressions. Across all A and τ combinations, with either 0.20a.u. or 0.15a.u. common input, R², slope, and intercept showed no consistent pattern (<u>Fig. S5A</u>). We then chose the LM_sim_ that best explained the experimentally observed dependency of inhibition on firing rate for each of the 8 subjects, as previously done for the CSP simulations (<u>Fig. S2D</u> and <u>S5C</u>; <u>table S5</u>).

**Fig. 5.**
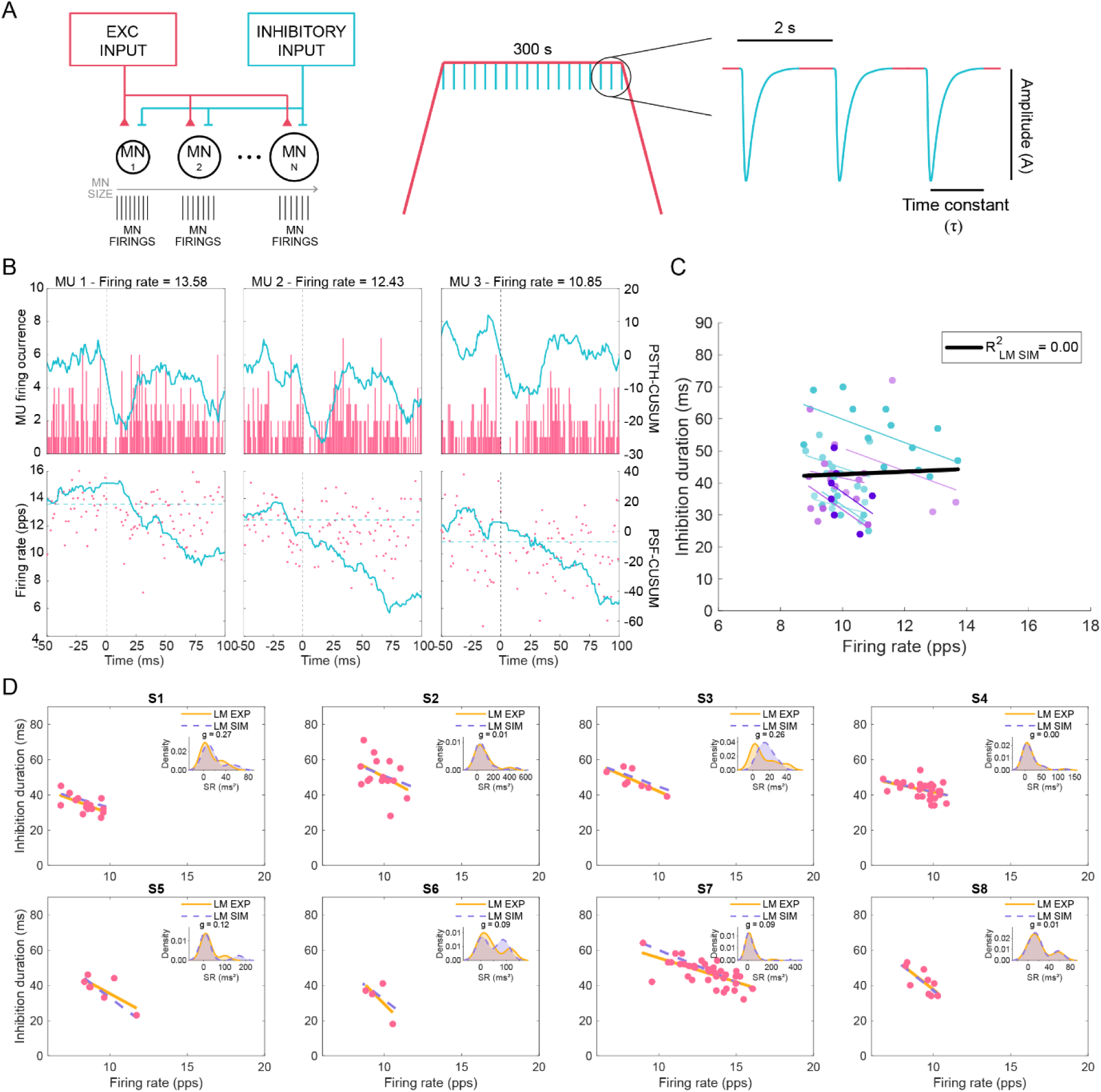
Characterization of reciprocal inhibition on individual motoneurons with computational modelling. **(A)** Schematic of the *in silico* computational model used to simulate the common input and inhibitory input delivered to motoneurons (MN) of varying size (left), during artificial voluntary contraction with periodic inhibitory inputs to motoneurons delivered during the plateau phase (right). **B)** Examples of PSTH (top) and PSF (bottom) of simulated MUs at different frequencies, with cumulative summation (CUSUM) traces in blue. **(C)** Simulated datapoints and respective linear fits for inhibition duration relationship with discharge rate across the LM_sim_ optimized for each subject (different shades of blue per optimized LM_sim_); black line represents the linear fit for the simulated group data with respective R^2^ shown. **(D)** Experimental MU inhibition durations and firing rate (pink dots) with respective regression line (orange line; same LM_exp_ as in <u>Fig. 4E</u>) and optimal LM_sim_ fit for each participant (dotted purple line). Inset plots show distributions of square of residuals (SR) as kernel density estimates and the corresponding Hedges’ *g* value; pps – pulse per second.

When comparing the optimized LM_sim_ for each subject (<u>Fig. 5C</u>) and experimental fits (LM_exp_; <u>Fig. 5D</u>), by analysing the distributions of square of residuals obtained from experimental data points against linear fits, we observed small effect sizes for subjects 1 and 3, while no effects were found for the other participants. Although the LM_sim_ realizations did not exhibit a clear pattern and the selection process appeared almost random (<u>Fig. S5C-D</u>), they still managed to fit experimental reciprocal inhibition data reasonably well. These results highlight the variability in the success of our *in silico* model in capturing experimentally observed dependencies of inhibition duration on firing rate. While these biophysical models generated a range of fits that happened to align with the experimental data, the weak R² values suggest that this agreement may be more coincidental than indicative of a strong underlying relationship.

## DISCUSSION

In this work we used HDsEMG to study the temporal properties of the CSP in a hand muscle (FDI) and reciprocal Ia inhibition in the lower leg (TA muscle). We successfully sampled multiple MUs per individual and demonstrated that functional inhibition duration is best interpreted in the context of varying MU discharge rates, emphasizing the need for subject-specific analysis. By measuring inhibition using firing rates, we are not measuring inhibitory postsynaptic potentials (IPSP), but rather how the inhibition affects motoneuron output. For example, large amplitude IPSPs may appear to be of shorter duration if they recruit active currents (e.g. hyperpolarization-activated current [I_h_] or CaV3 currents), leading to earlier onset firing. We also used *in silico* modelling, which indicated that the underlying firing rates and characteristics of the inhibitory inputs rather than variations in S-type motoneuron size play roles in determining the functional duration of inhibition across different MUs.

### Subject-specific analysis of spinal inhibitory circuits

Inter-individual variability is intrinsic to the human nervous system, as evidenced by neuroimaging^47^, electroencephalography^47^, transcranial magnetic stimulation^48,49^ and electromyographic and kinematic studies^10,50,51^. Such variability significantly influences computational modelling of human neurophysiology^10,52–54^. In clinical neurophysiology, traditional methods such as sEMG or iEMG yield low-dimensional data due to their technical limitations, which limit the assessment of inter-individual differences. For example, the electrical response obtained with sEMG does not inform about individual MUs, and single MU iEMG is invasive leading to sampling of only a few units^9^. Such constraints result in data homogeneity being assumed *a priori*, leading to individual MU data being aggregated across subjects and analysed collectively. In contrast, with HDsEMG we were able to extract multiple units per individual, resulting in datasets with a hierarchical structure: that is, MU data is nested within subjects. This not only opens the possibility for clinically relevant subject-specific studies with HDsEMG, but also introduces more rigour in data interpretation by addressing data dependencies, thus reducing the risk of false positives^55–57^.

For example, recent work using iEMG established weak relationships (R^2^<0.33) between CSP inhibition duration and MU discharge rate (R^2^=0.30)^13^, similar to our findings when we plotted all sampled MUs together (R^2^=0.24). But by sampling an average of 7 units per individual, unlike for iEMG^13^, we were able to establish linear regressions for each subject, and employ more rigorous statistical approaches that account for the high (80%) inter-individual variability of the CSP. Altogether, our findings highlight the potential of HDsEMG in the study of inhibitory spinal circuits by permitting subject-specific clinical insights.

### CSP and reciprocal inhibition: methodological and physiological considerations

We observed that inter-participant variability, the strength of inhibition, and the relationships between discharge rate and inhibition duration, were not the same for the CSP measured from the FDI and reciprocal inhibition recorded from the TA muscle. These differences could be influenced by multiple factors such as different stimulation paradigms to evoke CSP and reciprocal inhibition^22^, synaptophysiology, upper and lower limb motoneurons, and microcircuit physiologies and contraction profiles.

For the CSP we used a stimulation protocol designed to robustly activate A-delta fibers mediating the withdrawal reflex^5,26–28^. To evoke reciprocal inhibition, we stimulated the tibial nerve at 1.1x the minimum H-reflex amplitude threshold to minimize activation of afferents other than the largest (e.g. proprioceptive Ia afferents), and crosstalk contamination from heteronymous monosynaptic Ia afferent excitation to the TA muscle^58,59^. Therefore our stimulation protocol likely recruited robust Ia reciprocal inhibition. The synaptic input received by motoneurons when evoking the CSP is considerably slower^60^ likely due to GABA_A_ receptor activation^61^, whereas reciprocal inhibition is purely glycinergic^62,63^; such factors could lead to differences between CSP and reciprocal inhibition in the strength and timing of inhibition in motoneurons^64,65^. Furthermore, although TA and FDI muscles have similar muscle fibre composition, contractile properties and discharge rates^66^, physiological differences may exist between upper and lower limb circuitries that may affect the strength of the correlations with discharge rate. One must also consider the impact of active conductances in our PSTH-PSF estimations, for example, computational studies have suggested that activation of I_h_ at more hyperpolarized voltages, which may have been elicited at stronger stimulation intensities may produce a faster rebound that could shorten inhibition^67^. Finally, one must consider that synaptic noise introduces variability in spike timing, which can disproportionately affect PSF and PSTH estimates, particularly when detecting smaller-size functional inhibition^22,68^.

### Accuracy of *in silico* modelling of inhibitory spinal circuits

Biophysical simulations for the CSP revealed that R^2^ increased with higher inhibition amplitude and longer τ. More interestingly, the optimized parameters for each participant varied widely across subjects (<u>Fig. 2D</u> and <u>Fig. S3B</u>). Some individuals, particularly those near the limits of the hyperparameter space, showed poor fits (e.g. subject 1) while others had a better fit to the experimental HDsEMG data (e.g. subject 5). In some cases, multiple realizations could perform similarly (subject 2) and for others there were fewer realizations with low MSE (subject 6; <u>Fig. 2D</u> and <u>Fig. S3B-C</u>). Altogether these results highlight the complexity of simulating biological relationships for spinal microcircuits with robust inhibition.

Caution must be taken about the interpretation of the robustness and predictive power of the *in silico* models. The HDsEMG data for reciprocal inhibition did not yield any strong R² coefficients for the tested subjects (<u>Fig. 4D</u>), and simulating data reproducing experimental relationships with weak R^2^ values is inherently more challenging^69^. The optimized parameters for most subjects (6 out of 8) were located near the boundaries of the explored hyperparameter space, both near the lower and upper ends of A and τ (<u>Fig. S5C-D</u>), suggesting that our simulations may not have fully captured the range of physiological variability. While simulations were more realistic and performed well for circuits with robust inhibition such as the CSP (Fig. <u>S3A</u>), they were less reliable for pathways with shorter inhibition such as reciprocal inhibition. (Fig. <u>S5A-B</u>). In these cases, multiple factors, such as synaptic noise or fewer inhibited spikes, may influence the linear relationships between inhibition duration and MU discharge rate, potentially leading to unrealistic and oversimplified fits that happened to align with the experimental data^70^. These findings suggest that reciprocal inhibition measured in our experimental conditions, is not only physiologically weaker than CSP, but also more difficult to quantify and model accurately, thus stressing the need for caution when working with weak or noisy fits in both experimental and simulated HDsEMG data.

Motoneurons are complex neuronal cells characterized by an extensive dendritic tree, large soma, complex axonal excitability dynamics, high density and variety of synaptic inputs, and diverse, spatially localized ion channels and metabotropic receptors^71^. Such complexity would be best captured with multi-compartment models, but for our purposes we employed a simple two-compartment model as it has been shown that both single- and two-compartment models can successfully replicate variations in MU firing under different modulatory and synaptic inputs^10,72–74^. Our model employed simplified ionic conductance parameters, omitting additional active conductances (e.g. persistent inward currents) that could potentially influence the simulated MU firing rates. Despite these constraints, our *in silico* simulations successfully reproduced the observed MU firing rate-inhibition duration relationships across most subjects, indicating that synaptic input dynamics primarily drive these phenomena.

In line with recent work^10^, we emphasize the importance of subject-specific parameter considerations when generating computational models for the study of spinal circuits. Biophysical models accounting for individual subjects not only better replicate experimental findings, but are also essential for understanding spinal connectivity and functional impact of synaptic inputs. This is because direct, invasive measurements are not feasible in humans^70^, thus making *in silico* models indispensable for studying human spinal synaptophysiology.

### Dependence of inhibition duration on firing rate and motoneuron size

Our computational model was fine-tuned on experimental data obtained at 10% MVC, with simulations reflecting the behaviour of early-recruited MU, likely representing a portion of small-sized motoneurons within the FDI muscle (the “size principle”^35,41^). The differences in soma capacitance between the largest and smallest motoneurons in our simulated dataset did not exceed 15%. However, incorporating parameters from fast motoneurons typically recruited at higher MVCs could reveal ∼2-fold differences in soma capacitance between the smallest and largest motoneurons^72^ (<u>Fig. 3</u>). Simulations restricted to small motoneurons may risk overestimating the uniformity of inhibition duration across the motor pool, as they may not represent true neuromuscular physiology. Of note, though, smaller motoneurons would be more susceptible to an incoming inhibitory input than their larger counterparts^41,45^, implying that any potential effects of motoneuron size on inhibition duration would likely be more readily detectable in our simulations.

Interestingly, linear increases in common input revealed greater rates of change in firing rate amongst earlier recruited units with later-recruited units plateauing in their discharge rates at higher contraction levels^75^. Furthermore, the FDI stops recruiting units at lower MVCs when compared to other muscles^75^, suggesting that our simulations replicating an experimentally observed 10% MVC may have included enough MU diversity to detect any size-dependent effects on inhibition duration with varying firing rate.

While we cannot exclude the possibility that alternative parameters outside the explored hyperparameter space could produce different outcomes, we tested optimized parameters from two subjects with markedly different values of A (1.8 and 2.8 a.u.) and τ (7 ms and 19 ms). In both cases, motoneuron size had only a minor effect on inhibition duration, suggesting that the strength of inhibition is unlikely to play a role in the observed relationship.

## Conclusion

Our study highlights HDsEMG as a tool that offers unparalleled resolution for subject-specific studies of spinal inhibitory microcircuits. By integrating experimental data with *in silico* modelling we further extend the scope of HDsEMG in elucidating both circuit function and the biophysical properties of its neuronal elements. We hope that our work encourages broader adoption of HDsEMG as an approach to probe motor circuit function in health and disease, given that its technical advantages open new frontiers in the quality and depth of data attainable from human subjects.

## MATERIALS AND METHODS

### Participants

18 healthy subjects (10 male, 8 female, aged 25±5 years, height 172±10cm) were recruited for this study. The study was approved by Imperial College London ethics committee (reference number 18IC4685) and the experiments were conducted in accordance with the Declaration of Helsinki and all the participants signed a written consent form prior to the experiments.

### Procedures

#### High-density surface electromyography

Prior to the application of the HDsEMG electrodes, the skin was prepared by applying abrasive paste and water. For the CSP experiment, one HDsEMG grid (64 electrodes,13 rows and 5 columns; 1mm electrode diameter, 4mm inter-electrode distance; OT Bioelettronica, Italy) was placed on the FDI of the dominant hand. Two wet wristbands were used as reference and grounding electrodes on the right wrist of the participants. For the reciprocal inhibition experiment, a customized HDsEMG grid (256 electrodes, 32 rows and 4 columns; 0.5mm electrode diameter, 4 mm inter-electrode distance, OT Bioelettronica, Italy) was placed on the TA of the dominant leg. Two HDsEMG grids (128 electrodes, 26 rows and 5 columns; 0.5mm electrode diameter, 4mm inter-electrode distance, OT Bioelettronica, Italy) were placed on the medial and lateral head of the soleus (SOL). Two wet wristbands were used as reference and grounding electrodes on the right ankle of the participants. All the signals from the HDsEMG electrodes were acquired at 2042.48Hz in monopolar mode (Quattrocento, OT Bioelettronica, Italy).

#### Cutaneous Silent Period

To study CSP, subjects were asked to perform a pinching task between the medial side of the third phalanx of the first digit of the hand and the thumb finger while stimulation on the fifth digit of the hand was delivered. Subjects were sat on a chair while conducting the experiment. A load cell (FC22, Measurement Specialties, US) was used to measure the force applied on the index finger. The force signals were acquired at 2042.48Hz simultaneously with the HDsEMG and displayed as visual feedback. Subjects were asked to perform their maximum voluntary contraction (MVC) for the pinching task. Then, stimuli were delivered using two stainless steel rings which were placed between the first and the second phalanges of the fifth digit of the hand. Stimulation pulse width was set to 200µs and conductive gel was applied between the ring electrodes and the skin to reduce impedance - extra care was taken to remove any conductive paste between the ring electrodes to prevent shorting. Sensory threshold (ST), defined as the minimum stimulation intensity that generated a subjective perception on the finger, was found for each subject and stimulus intensity was set at 10 times this ST for each subject. After familiarization with the execution of the force-tracking task, subjects were asked to perform a submaximal trapezoidal isometric contraction at 10% MVC (2s ramp up, 200s plateau, 2s ramp down) while single electrical stimuli were delivered on the fifth digit of the hand (1.8±0.2s inter-stimulus interval; ∼111 stimuli) during the plateau.

#### Reciprocal inhibition

To study reciprocal inhibition on the TA, subjects were asked to perform an isometric dorsiflexion task while electrical stimuli were delivered on the tibial nerve. Subjects were sat on a chair while their leg was fixed into an ankle dynamometer (OT Bioelettronica, Italy) using straps (ankle position at 10°, 0° being the neutral foot perpendicular to the shank; knee at 75°). The dorsiflexion force signals were acquired at 2042.48Hz simultaneously with the HDsEMG and displayed as visual feedback. Subjects were asked to perform their MVC for the dorsiflexion task. A stimulation electrode (7.5×13cm adhesive electrode, ValueTrode, Axelgaard, Denmark) was placed as anode over the patella, while a stimulation electrode was placed as cathode (1cm diameter customized metal ball) over the middle part of the popliteal fossa. The final stimulation position of the cathode was optimised for each participant to elicit a clear H-reflex response in the SOL^76^. Stimulation pulse width was set to 1ms and conductive gel was applied to the cathode to reduce skin impedance and discomfort. The stimulation intensity was selected as 1.1x of the stimulation intensity necessary to elicit the minimum detectable H-reflex on the SOL (based on selection of monopolar channels from the medial head of the SOL). After familiarization with the performance of the force-tracking task, subjects were asked to perform a submaximal trapezoidal dorsiflexion contraction at 10% MVC (2s ramp up, 300s plateau, 2s ramp down) while single electrical stimuli were delivered to the tibial nerve (2.0±0.2s inter-stimulus interval; ∼150 stimuli) during the plateau. This stimulus rate was appropriate to avoid exacerbating post-activation depression^77^

#### *In silico* modelling

To model CSP and reciprocal inhibition circuits in the motor pool, we implemented a Hodgkin-Huxley model with dendritic and somatic compartments^72^. The differential equations to compute the membrane voltage for each motoneuron was computed using Equation 1 and solved with the fourth-order Runge-Kutta method with a 0.2ms step resolution^78,79^.

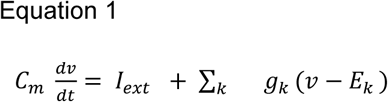

Where *C_m_* represents the membrane capacitance, and *I_ext_* is the external synaptic input to the cell. The conductance of a specific synaptic or intrinsic ion channel *k* is denoted by *g_k_*, while *v* represents the compartment’s voltage. The term *E_k_* corresponds to the reversal potential associated with each conductance, which may represent either the equilibrium potential of a specific ion in the case of intrinsic currents, or the effective reversal potential for excitatory and inhibitory synaptic inputs. Since each compartment has distinct ion channels and inputs, their governing equations differ. The somatic compartment includes sodium channels and both fast and slow potassium channels, while the dendritic compartment contains sodium channels along with excitatory and inhibitory synaptic inputs (Equation 2).

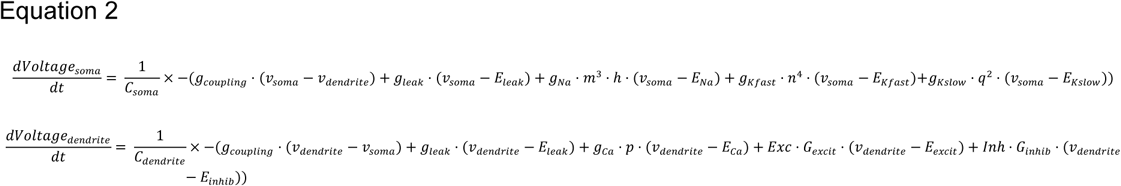

With *C_soma_* and *C_dendrite_* representing the capacitance of each compartment; *g_coupling_* the electrical coupling between somatic and dendritic compartments - computed using axial resistivity and compartment geometry, modelling the soma and dendrite as cylinders; *v*_soma_ and *v*_dendrite_ are the voltages at a given time *t* for each motoneuron and compartment; *g_leak_* are the conductances for leak currents; *g_Na_* represents voltage-gated Na^+^ channels; *g_Kfast_* and *g_Kslow_* indicate fast and slow-kinetics voltage-gated K^+^ channels, respectively.

Gating variables followed first-order kinetic equations, representing the probability of ion channels being open or closed; *m* and *h* govern the activation and inactivation of sodium channels, *n* and *q* control the activation of fast and slow potassium channels, respectively, and *p* regulates calcium channel activation; E_Na_, E_K,_ E_Ca_ and E_leak_ are the equilibrium potentials for Na^+^, K^+^, Ca^+^ and leak channels; E_excit_ and E_inhib_ are the reversal potentials for excitatory and inhibitory inputs; *Exc* represents the excitatory common input (a.u.) given to the motoneuron pool – in this case a trapezoid signal – and *Inh* (a.u.) is the inhibitory input received reproducing the CSP or reciprocal inhibition; and G_excit_ and G_inhb_ are the synaptic conductance for excitation and inhibition. To introduce biological variability, each motoneuron receives an excitatory input and a white Gaussian independent noise that follows a distribution of N (0,0.25σ^2^). Motoneuron firing occurred when somatic voltage exceeds a threshold calculated as the product of the minimum current to elicit an action potential and input resistance.

The motoneuron parameters were obtained from *in vivo* motoneuron recordings obtained from S-type motoneurons from decerebrated cats^80–85^, and linearly interpolated across *n* simulated motoneurons, ranging from the smallest to the largest motoneuron sizes (see <u>table S6</u>). Particularly, soma and dendrite length and diameter were linearly varied within the range of parameters for the *n* simulated motoneurons, using these values to estimate the soma and dendrite conductance and capacitance for each neuron. S-type motoneurons were specifically chosen as they best represent the motoneuron pool recruited at low contraction intensities (10% MVC) during experimental HDsEMG recordings.

A motor pool of 20 motoneurons was simulated, each receiving trapezoidal common synaptic input mimicking the force trajectory performed in the experimental conditions: a 2s ramp-up and ramp-down phase with a 200s plateau phase in between for the CSP modelling, and 300s plateau for the reciprocal inhibition modelling. Prior to the modelling of the inhibitory responses, we adjusted the amplitude of the excitatory common input (between 0.15 to 0.3a.u.) in order to replicate the firing rates measured with HDsEMG from the FDI and TA muscles. We compared the experimental firing rate – averaged across all participants - and checked that it closely mimicked the simulated data by computing the error between them (error∼0) and also by implementing a Kolmogorov-Smirnov test. This test confirmed that both datasets followed the same distribution (p=0.48). After testing different amplitudes for the excitatory common input, we found that an amplitude of 0.2a.u. replicated the experimental average firing rate well for both muscles. Additionally, for reciprocal inhibition, an amplitude of 0.15a.u. for the common input was simulated to minimize noise effect of background firing rate on estimating inhibition duration, without deviating from experimentally observed firing rates (<u>Fig. S5</u>).

The inhibitory input (*Inh*) replicating the inhibitory responses elicited through electrical stimulation was modelled as an alpha function (a.u.) with time constant (τ) and amplitude (A). This approach was chosen to capture the rapid onset of inhibition followed by a gradual decay (see <u>Fig. 2A</u> and <u>Fig. 5A</u>).

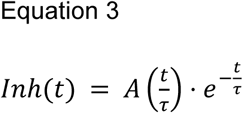

To reproduce the inhibitory responses observed in the experimental CSP and reciprocal inhibition, we conducted a hyperparameter sweep to explore different combinations of inhibition amplitude (A) and time constant (τ). This allowed us to map a parameter space that could generate motoneuron inhibition profiles consistent with the experimental data. The inhibitory input (*Inh*) to the simulated motoneurons was delivered every 1.8s for CSP (<u>Fig. 2A</u>) and every 2s for reciprocal inhibition (<u>Fig. 5A</u>). Inhibition amplitudes (A) ranged from 1 to 3a.u., varying in 0.2a.u. increments, while the time constant (τ) varied between 7 and 20ms for CSP and 1-4ms for reciprocal inhibition in 1ms steps. The resulting motoneuron spike trains were downsampled from 20KHz to 2kHz to match the experimental MU spike trains and the inhibitory responses were analysed manually.

To investigate the impact of motoneuron biophysical properties, namely motoneuron size, on the duration of inhibition for CSP, we simulated a larger pool of 40 motoneurons using the previously described paradigm. We doubled the number of motoneurons for these simulations to achieve a higher resolution in mapping the intrinsic property gradients of motoneurons. In this case, the inhibitory input strength was held constant within each simulated motoneuron pool, triggered every 1.8s with A and τ values were selected so they closely matched experimental data from one of the subjects (subject 4, randomly selected; <u>Fig. 3A</u>). Meanwhile, for each motoneuron pool simulated the common excitatory synaptic input was varied (*Exc* varying from 0.21a.u. to 0.28a.u., with 0.01a.u. increments). The output motoneuron spike trains were downsampled to 2kHz and the inhibitory responses were analysed manually following the methods described below. The goal of these simulations was to generate motoneurons with different sizes that, despite receiving the same inhibitory drive, discharged at similar firing rates due to differences in the strength of the common excitatory input.

### Data analysis

#### HDsEMG decomposition

Monopolar HDsEMG signals were decomposed into individual MU spiking activity using a validated blind source separation algorithm^86–88^. This algorithm provides a solution to the inverse EMG convolutive mixing model by estimating the separation vectors (MU filters) that yield the sources of the EMG signals (individual MU spike trains). Prior to the application of the decomposition algorithm, signals were band-passed (20-500Hz, second order Butterworth) and visually inspected to reject channels with low signal-to-noise ratio. Since the stimulation artifact and the presence of evoked compound action potentials could bias the identification of MUs, the signals contained in the interval 5-100ms relative to the stimuli where not inputted into the decomposition algorithm. Due to the extended length of the recordings, a window of 30s selected from the plateau of the isometric contraction was selected to extract the MU filters. Then, the MU filters were extended to the whole recording and the MUs identified in the 30s plateau were tracked throughout the entire trial. All HDsEMG signals were decomposed using the same procedure.

Among all the MUs detected, only those with pulse-to-noise ratio ≥ 28 dB^89^, coefficient of variation (CoV) of their discharge rate <0.3 and stable discharge patterns were retained for analysis. Despite the isometric contraction task yields a stable MU discharge rate, the responses evoked via electrical stimulation could elicit alterations on the discharge rate (inhibition or excitation). Traditional manual edition of MU spike trains and refinement of MU filters could bias in the edition of the inhibitory/excitatory responses. Thus, the MU spike trains output from the decomposition algorithm did not undergo manual edition in this study. MU spike trains and HDsEMG signals were downsampled to 2kHz for the subsequent analysis.

To estimate the onset of the motor unit action potential (MUAP) relative to the discharge times identified by the decomposition algorithm, MUAPs were calculated by spike-averaging all the HDsEMG channels, and the channel with the highest peak-to-peak amplitude was selected to estimate the MUAP onset time^90^. Using this channel, a threshold set at 20% of the maximum amplitude of the MUAP was set to define the onset time of the MUAP, which was checked visually for all the MUs included in the analysis.

#### Inhibitory responses

Instead of manual edition, an automatic procedure was applied to remove outliers in the discharge pattern by removing instantaneous discharges above the median discharge rate plus 2.5x the standard deviation and below 50% of the median discharge rate. The resultant MU spike trains were used to estimate the inhibitory responses with the peri-stimulus time histogram (PSTH) used to estimate the latency (i.e. start of inhibition) and the peri-stimulus frequencygram (PSF) for duration (i.e. end of inhibition) as previously described^6,7,20^. In summary, the PSTH, representing the number of MU firing occurrences locked around the stimulation, and the PSF, representing the instantaneous discharge rates, were computed in bin widths of 1ms. The cumulative sums (CUSUM) of both PSTH and PSF were calculated. The reasoning for using both PSTH and PSF is the following: PSTH reflects the probability of MU firing relative to the stimulus, thus making it suitable for determining inhibition onset. However, it cannot capture the full duration of inhibition since it relays information from changes in firing probability. PSF tracks instantaneous discharge rate throughout time, and since inhibition affects the firing rate rather than completely silencing all the units especially toward the rise phase of the IPSP (i.e. late, weaker phase), it provides a continuous measure of how long the synaptic inhibition persists. So by using PSTH for onset and PSF for end of inhibition, we ensure a more precise characterization of the inhibition duration^6,7,20^. Since the stimulation artifact and the presence of direct motor response (M-wave) could be merged in the MU spike train leading to inaccurate discharge identification, the CUSUM was not updated in the window comprising ±20ms around the stimulation. Inhibition events were identified visually and set as genuine inhibitory responses when the amplitude between two inflexion points in the PSTH-CUSUM was greater than the maximum variation in the pre-stimulus window (−200ms, -30ms, limits <u>Fig. S1</u>). Genuine inhibitory events were detected according to the criteria above in ∼80% and ∼70% of all MUs sampled for CSP and reciprocal inhibition, respectively. The motoneurons which have a weak inhibition but not significant (i.e. inhibition size smaller than the CUSUM limits) were not included into linear model generation, thus resulting in variable number of motoneurons in each realisation for modelling. The inhibition latency and end time points were selected manually in the PSTH-CUSUM and PSF-CUSUM as the first clear deflection point of a trough, and the last deflection point or peak of the trough, respectively. The inhibition duration was computed as the time interval between the inhibition latency estimated in the PSTH and the inhibition endpoint selected in the PSF. The inhibition amplitude was computed as the difference between the CUSUM values at the end and onset of the inhibition time points for both PSTH and PSF, and reported in the supplemental material (tables <u>S7</u>-S10).

To estimate the inhibitory responses in the global EMG, raw sEMG signals were digitally band-pass filtered (20-500Hz, second order Butterworth) and then rectified and segmented in 600ms windows around the stimuli. The rectified sEMG signals were relativized to the baseline activity, which was calculated as the averaged rectified sEMG in the time interval from 200 to 25ms prior the stimulus. The start of inhibition was determined manually as the time when the signal was lower than baseline for at least 5ms, while end of the inhibition was manually set as the time when the signal was no longer lower than baseline for at least 5ms. Inhibition duration was estimated as the time difference between the start and the end points (see <u>Fig. S1</u>).

Since inhibition measured through HDsEMG during voluntary contractions is influenced by discharge rate, parameters were interpreted in relation to MU firing frequencies. The experimental inhibitory responses (firing rate and inhibition duration) were fitted to a linear model (LM_exp_), and to evaluate how discharge rate explained variations in inhibition duration, we relied on the coefficient of determination (R^2^) metric as a measure of goodness-of-fit^30,31^. This R^2^ was calculated as:

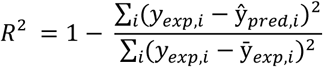

where y_exp,i_ and ȳ_exp,i_ is the experimental datapoint and experimental mean for the i^th^ observation, respectively, and ŷ_pred_ is the predicted inhibition duration from the linear fit (Fig. <u>S2A</u>). Although the relationship between inhibition duration and MU firing rate is the ideal proxy for estimating inhibition, we still report in the supplemental information pertaining to PSTH and PSF amplitudes (tables <u>S7</u>-<u>S10</u>).

For all *in silico* simulations, the inhibition response for each motoneuron was manually quantified using PSTH and PSF methods, following the same procedure as for the experimental data analysis described above (<u>Fig. 2B</u>). Then, the inhibitory responses (firing rate and inhibition duration) obtained from each realisation of the simulation were fitted to a linear model (LM_sim_; Fig. <u>S3A</u> and <u>S5A-B</u>). To select the optimal LM_sim_ for each subject, we first inputted their experimental firing rate into all the candidate LM_sim_ obtained from all the realizations of the biophysical model across the different combinations of parameters A and τ. The selection process prioritized minimizing the Mean Squared Error (MSE), which has the same numerator as the R^2^, ensuring that the chosen LM_sim_ provided the best prediction of inhibition duration across subjects by reducing overall deviation from experimental data while preserving the underlying relationship between firing rate and inhibition duration (see <u>Fig. S2D</u>):

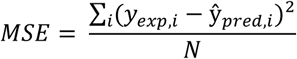

Inhibition duration, PSTH and PSF amplitudes for CSP and reciprocal inhibition for the simulated data pertaining to the LM_sim_ selected for each subject are reported in supplemental section (tables <u>S11</u>-<u>S16</u>).

For the *in silico* simulations exploring the impact of motoneuron biophysical properties on inhibition duration, the resulting motoneurons across all the simulations were clustered into 3 main groups based on their firing rate (11-12Hz, 13-14Hz, 14-15Hz). This clustering permitted to evaluate how cell size (i.e. capacitance) could explain variability in inhibition duration within fixed MU discharge rate intervals (see <u>Fig. 3B</u>).

#### Statistical analysis

Statistical analyses, computational simulations, plots and figures were generated using MATLAB R2022b (Mathworks, USA) and Microsoft Excel version 2208 (Microsoft, USA). The main statistics focused on relationships between motor unit activity variables, including:

1. Inhibition duration and firing rate;
2. PSTH inhibition amplitude and firing rate;
3. PSF inhibition amplitude and firing rate;
4. Motoneuron capacitance and inhibition duration.

For each of the relationships we performed two different complementary analyses:

1. Grouped analysis: here we considered all the MUs as independent, and pooled experimental MU data from all subjects (<u>Fig. 1D</u> and <u>Fig. 4D</u>), or simulated MUs pertaining to the optimized LM_sim_ for each subject (<u>Fig. 2C</u> and <u>Fig. 5C</u>), and fitted a linear regression model (LM) using the *fitlm* function in MATLAB^91^. The coefficient of determination (R^2^) was used to infer the strength of the linear regression in explaining the variance in each relationship and therefore the LM’s ability to capture meaningful trends within the pooled data^92^. For better interpretation of R^2^ values obtained, we will consider the threshold values of 0.67, 0.33, or 0.19 as strong, moderate and weak coefficients^32,33^ (<u>Fig. S2</u>). Additionally, to assess the statistical significance of the linear fit, we report p-values obtained from *fitlm*^91^. These p-values are computed from the t-statistic (*t*), which evaluates whether the slope (β) of the regression line significantly differs from zero^93^:

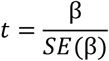

where *SE* is the standard error of the slope. The corresponding p-value is derived from a t-distribution with degrees of freedom (n−2), with statistical significance set at 0.05^93^.

1. Subject-specific analysis: we analysed the data by taking into consideration inter-group variabilities. Here we fitted a linear regression model for the experimental data pertaining to each subject (LM_exp_; <u>Fig. 1E</u> and <u>Fig. 4E</u>), or to the simulated MUs for each realization of the in silico biophysical model (Fig. <u>S3</u> and <u>S5</u>), generating linear regression models for the simulated data (LM_sim_). Only realizations with more than 2 MUs with detectable inhibition were fitted. For the experimental data (<u>Fig. 1E</u> and <u>Fig. 4E</u>), and the simulated MU data pertaining to the optimized LM_sim_ selected for each subject (tables <u>S11</u>- <u>S16</u>), we also fitted a random intercept (Model 1) and a random intercept and random slope (Model 2) linear mixed model (LMM) using the *fitlme* function from MATLAB under the restricted maximum likelihood fitting method^94^ as follows:

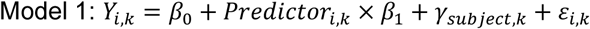

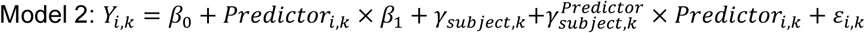

With Y_i,k_ representing the collected datapoint for “Inhibition duration”, “PSTH inhibition amplitude” or “PSF inhibition amplitude”, belonging to the i^th^ observation obtained from the k^th^ subject; β_0_ is the fixed intercept; Predictor_i,k_ is the predictor variable (“Firing rate”) for observation i within k and β_1_ is its corresponding coefficient ; γ_subject,k_ represents the random effect intercept for the k^th^ subject, in order to capture subject-specific deviations from the fixed intercept; γ^*Predictor*^_*subject,k*_ is the random effect slope for the predictor variable in the k^th^ subject, in order to account for subject-specific variability in the predictor and outcome relationships (e.g. slope for inhibition duration *vs* firing rate is adjusted for each subject according to their firing rate); ε_i,k_ is the residual error for observation i in subject k.

From the *fitlme* output, we reported the predicted value for the intercept and the estimated difference for the fixed effects along with respective 95% confidence intervals (CI). The *fitlme* also provides the standard error and its 95% CI for each of the random effects, which we used to estimate the intraclass correlation coefficient (ICC). The ICC quantifies the proportion of total variance attributable to between-subjects variability, thus relaying information into the sources of variability in our hierarchical datasets^57,95^. For Model 2, we also report the correlation and 95% CI for the random slope and intercept interaction, noting that overfitting sometimes led to singular fits (“Singular”; see tables), indicated by a correlation of ±1. We also report the R^2^, a metric used to gauge the variance explained by the LMM^96^.

For graphical representation and data interpretation, we selected either LMM Model 1 or Model 2 based on a LMM comparison. We used the *compare* function in MATLAB^97^, to perform a likelihood ratio test, which compares each of the LMM’s log-likelihood, Akaike information criterion (AIC), Bayesian information criterion (BIC) and provides a p-value of the test. These metrics evaluate the balance between model fit and complexity^98^, with the LMM exhibiting the lowest AIC/BIC, highest log-likelihood, and a p-value lower than 0.05 from the likelihood ratio test statistic being selected.

To select the optimized LM_sim_ for each subject, we compared the R^2^ experimentally determined with recorded data (LM_exp_; <u>Fig. 1E</u> and <u>4E</u>) with the R^2^ of the all the generated LMs_sim_ predicted with experimental firing rates (Fig. <u>S2-S5</u>; <u>Fig. 2D</u> and 5D). The R^2^ metric is a common goodness-of-fit validation metric for biophysical models^69^, as it quantifies the proportion of variance in experimental data captured by the simulations, thus allowing to assess how well our *in silico* models aligned with the observed experimental trends for each subject^69,99^. For the simulations with varying motoneuron size (<u>Fig.3</u> and <u>S4</u>), data obtained were grouped into three different firing frequency groups (11-12Hz, 12-13Hz, and 13-14Hz) with varying motoneuron (MN) capacitance. A random intercept LMM was used to compare the changes in inhibition duration:

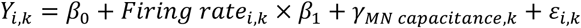

With *Y_i,k_* representing “Inhibition duration” for the *i^th^*observation obtained for the *k^th^* motoneuron capacitance value; *β_0_* is the fixed intercept; *Firing rate_i,k_* is the firing rate group belonging to the observation *i* within *k* with *β_1_* being its coefficient; *γ_MN capacitance,k_* represents the random effect term for the k^th^ motoneuron capacitance value; and *ε_i,k_* is the residual error for observation *i* in motoneuron group *k*. Additionally, to further understand the variability introduce by motoneuron size in predicting inhibition duration, we calculated the ICC for each firing rate group through a one-way random effects ICC model (1,1)^95,100^,which was estimated through the mean squares (MS) obtained from an one-way analysis of variance (ANOVA) performed with the *anova* function in MATLAB^101^:

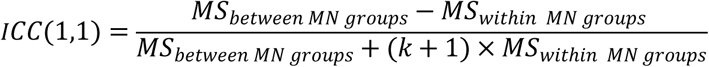

To provide a comparative metric when illustrating the optimized LM_sim_ and LM_exp_ for CSP and reciprocal inhibition (<u>Fig. 2D</u> and <u>5D</u>), we utilized Hedges’ *g* to compare the distributions of squared residuals (SR) obtained from experimental data points against those from linear regressions, as follows:

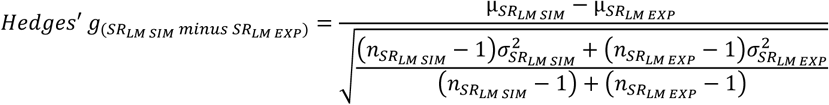

Hedges’ *g* can be used to calculate the effect size of the difference in residuals, with guidelines referring to small, medium and large effects as 0.20, 0.50 and 0.80, respectively^102^.

Data plots comparing inhibition duration estimated with the rectified EMG and PSF methods are shown as estimations plots. These plots display the minimum, first quartile, median, third quartile, and maximum value, with datapoints color-coded by subject. Additionally, bootstrapped (10,000 replicas) mean difference and Hedges’ *g* distributions (Kernell smooth) are shown, along with the mean (dot) and 95% confidence interval (CI; whiskers). The ‘0’ value is aligned with the mean difference of the *Rectified EMG* data, while the predicted mean difference for the *PSF* group is indicated by dotted horizontal lines. In the text, the values for inhibition duration pertaining to these estimation plots are reported as mean±standard deviation. The PSTH-CUSUM plots depict motor unit firing occurrence in bin widths of 1ms plotted against time with respective overlapped CUSUM trace, whereas the PSF-CUSUM plots illustrate individual MU firing events throughout time with respective CUSUM line. Correlation graphics are shown as scatter plots with all individual datapoints plus linear model and/or LMM regression lines. Inset plots depict distributions of squared residuals as kernel smooth density plots with Hedges’ *g* values positioned above. Hyperparameter heatmaps were generated to display linear fit parameters across the different A and τ combinations for each realization of the *in silico* simulations.

## Author contributions

Conceptualization, APV, FN, GO

Investigation, APV, FN, GO

Modelling Software, NTC, APV

Formal Analysis, APV, NTC, FN, GO

Writing – Original Draft, APV, NTC, FN, GO

Writing - Review & Editing - APV, NTC, MB, RMB, DF, FN, GO

Visualization - APV, NTC, FN, GO

Funding Acquisition, APV, NTC, MB, RMB, DF, FN, GO

Supervision, MB, RMB, DF, FN, GO

## Data and materials availability

### Lead contact

Further information and requests for resources and reagents should be directed to and will be fulfilled by the lead contact, Alejandro Pascual-Valdunciel.

### Materials availability

No new materials were created for this work.

### Data and code availability

All data and code that support this paper are made available through the Open Science Framework (https://osf.io/vutpx/)

---

**Table S1:** Fixed and random-effects variables and likelihood ratio test from random intercept (model 1) and random intercept and random slope (model 2) linear mixed models for the relationship between Inhibition duration and firing rate for cutaneous silent period (CSP) measured from first dorsal interosseous (FDI) muscle

|  | **Model 1:**<br>**Inhibition Duration ~ 1 + Firing rate + (1 \| Subject )** | **Model 2:**<br>**Inhibition Duration ~ 1 + Firing rate + (Firing rate \| Subject )** |
| --- | --- | --- |
| **Fixed effects** |  |  |
| (Intercept) | 304 ms [269, 338]<br>p = $6.76 \times 10^{-27}$ | 304 ms [244, 364]<br>p = $8.29 \times 10^{-15}$ |
| Firing rate | -15 ms [-17, -12]<br>p = $1.59 \times 10^{-17}$ | -15 ms [-20, -10]<br>p = $1.26 \times 10^{-7}$ |
| **Random effects** |  |  |
| Subject | 26 [16, 43], ICC = 0.60 | 80 [42, 152], ICC = 0.80 |
| Subject x Firing Rate | -- | 6 [3, 12], ICC = 0.06 |
| Correlation (Subject, Firing rate) | -- | -0.96 [-1, -0.77] |
| Residuals | 17 [14, 20], ICC = 0.40 | 14 [12, 17], ICC = 0.14 |
| **Model Fit and Comparison** |  |  |
| R<sup>2</sup> | 0.75 | 0.84 |
| AIC | 621.65 | 616.81 |
| BIC | 630.53 | 630.13 |
| Log-Likelihood | -306.83 | -302.4 |
| p-value | -- | 0.012 (Model 2 preferred) |

AIC: Akaike information criterion; BIC: Bayesian Information Criterion; ICC: Intraclass correlation coefficient

**Table S2.** Amplitude, time constant and common input values for the LM_sim_ selected from the CSP simulations for each of the subjects, with respective R^2^ from simulated MUs

| Subject | Excitatory input<br>(a.u.) | Inhibitory input<br>amplitude (a.u.) | Inhibitory input<br>time constant<br>(ms) | R<sup>2</sup> | p-value |
| --- | --- | --- | --- | --- | --- |
| 1 | 0.2 | 3 | 20 | 0.83 | 1.493E-05 |
| 2 | 0.2 | 2 | 17 | 0.89 | 6.99E-09 |
| 3 | 0.2 | 1.8 | 16 | 0.70 | 1.89 E-04 |
| 4 | 0.2 | 1.8 | 7 | 0.37 | 0.0269 |
| 5 | 0.2 | 2.8 | 20 | 0.78 | 6.26E-05 |
| 6 | 0.2 | 1.8 | 19 | 0.60 | 1.77 E-03 |
| 7 | 0.2 | 2.4 | 12 | 0.79 | 2.13E-05 |
| 8 | 0.2 | 2.8 | 19 | 0.92 | 2.37E-07 |
| 9 | 0.2 | 1.2 | 12 | 0.45 | 0.0116 |
| 10 | 0.2 | 1.4 | 18 | 0.72 | 1.23 E-04 |

**Table S3:** Intraclass correlation coefficient (ICC) and fixed and random-effects linear mixed model variables for inhibition duration at different firing rate groups with varying motoneuron capacitance for the simulated data from cutaneous silent period (CSP) from the first dorsal interosseous (FDI) muscle

|  | ICC capacitance |  |
| --- | --- | --- |
| Group | Optimized parameters subject 4 | Optimized parameters subject 8 |
| 11-12Hz | 0.16 | 0.16 |
| 12-13Hz | -0.24 | -0.23 |
| 13-14Hz | 0.06 | 0.02 |
|  | **Model:**<br>Inhibition Duration ~ 1 + Firing rate + (1 \| Capacitance) |  |
|  | Optimized parameters subject 4 | Optimized parameters subject 8 |
| **Fixed effects** |  |  |
| (Intercept)<br>11-12Hz | 109 ms [106, 111]<br>$p = 1.47 \times 10^{-120}$ | 99 ms [97, 102]<br>$p = 5.14 \times 10^{-123}$ |
| 12-13Hz | -12 ms [-16, -9]<br>$p = 4.00 \times 10^{-10}$ | -8 ms [-11, -5]<br>$p = 1.41 \times 10^{-6}$ |
| 13-14Hz | -18 ms [-22, -15]<br>$p = 3.23 \times 10^{-19}$ | -13 ms [-16, -10]<br>$p = 7.02 \times 10^{-14}$ |
| **Random effects** |  |  |
| Capacitance | 0 [,] ICC = 0.00 | 0.98 [0.01, 90] ICC = 0.12 |
| Residuals | 9 [8, 10] ICC = 1.00 | 7.41 [6.46, 8.50] ICC = 0.88 |
| **Model Fit** |  |  |
| R<sup>2</sup> | 0.42 | 0.33 |

Confidence interval for *Capacitance* random effect could not be accurately estimated.

**Table S4:** Fixed and random-effects variables and likelihood ratio test from random intercept (model 1) and random intercept and random slope (model 2) linear mixed models for the relationship between Inhibition Duration and firing rate covariation for reciprocal inhibition measured from tibialis anterior (TA) muscle

|  | **Model 1:**<br>**Inhibition Duration ~ 1 + Firing rate + (1 \|**<br>**Subject )** | **Model 2:**<br>**Inhibition Duration ~ 1 + Firing rate +**<br>**(Firing rate \| Subject )** |
| --- | --- | --- |
| **Fixed effects** |  |  |
| (Intercept) | 71 ms [61, 81]<br>p = $3.01 \times 10^{-26}$ | 74 ms [64, 83]<br>p = $8.05 \times 10^{-31}$ |
| Firing rate | -3 ms [-4, -2]<br>p = $1.27 \times 10^{-10}$ | -3 ms [-4, -2]<br>p = $1.15 \times 10^{-10}$ |
| **Random effects** |  |  |
| Subject | 9 [5, 15], ICC = 0.60 | 5 [1, 27], ICC = 0.44 |
| Subject x Firing<br>Rate | -- | 0.3 [0, 4], ICC = 0.03 |
| Correlation<br>(Subject, Firing<br>rate) | -- | 1 [Singular fit] |
| Residuals | 6 [5, 7], ICC = 0.40 | 6 [5, 6], ICC = 0.53 |
| **Model Fit and**<br>**Comparison** |  |  |
| R<sup>2</sup> | 0.56 | 0.56 |
| AIC | 800.63 | 803.83 |
| BIC | 811.78 | 820.55 |
| Log-Likelihood | -396.31 | -395.91 |
| p-value | -- | 0.669 (Model 1 preferred) |

AIC: Akaike information criterion; BIC: Bayesian Information Criterion; ICC: Intraclass correlation coefficient

**Table S5.** Amplitude, time constant and common input values for the LM_sim_ selected from the reciprocal inhibition simulations, with respective R^2^ and p value from simulated MUs

| Subject | Excitatory input<br>(a.u.) | Inhibitory input<br>amplitude (a.u.) | Inhibitory input<br>time constant<br>(ms) | R<sup>2</sup> | p-value |
| --- | --- | --- | --- | --- | --- |
| 1 | 0.15 | 1.2 | 5 | 0.33 | 0.42 |
| 2 | 0.2 | 1.6 | 2 | 0.08 | 0.71 |
| 3 | 0.15 | 2.6 | 4 | 0.12 | 0.29 |
| 4 | 0.15 | 2.4 | 4 | 0.02 | 0.71 |
| 5 | 0.15 | 1.4 | 2 | 0.45 | 0.10 |
| 6 | 0.15 | 2 | 2 | 0.78 | 0.31 |
| 7 | 0.2 | 2.4 | 3 | 0.35 | 0.03 |
| 8 | 0.15 | 1.8 | 3 | 0.18 | 0.35 |

**Table S6.** Parameter values adopted in the *in silico* modelling of S-type motoneurons.

| Variable | Definition | Value |
| --- | --- | --- |
| $l_D$ | Dendrite length | 5.50–6.80mm |
| $r_D$ | Dendrite radius | 20.75–31.25μm |
| $l_S$ | Soma length | 77.50–82.50μm |
| $r_S$ | Soma radius | 38.75–41.25μm |
| $C_m$ | Membrane specific capacitance | 1μF/cm<sup>2</sup> |
| $g_{Na}$ | Maximum sodium channel conductance | 30mS/cm<sup>2</sup> |
| $g_{Kf}$ | Maximum fast potassium channel conductance | 4mS/cm<sup>2</sup> |
| $g_{Ks}$ | Maximum slow potassium channel conductance | 16-25mS/cm<sup>2</sup> |
| $g_{Ca}$ | Maximum calcium channel conductance | 0.038-0.029 mS/cm<sup>2</sup> |
| $E_{Na}$ | Sodium reversal potential channels | 120 mV |
| $E_K$ | Potassium reversal potential channels | -10mV |
| $E_{Ca}$ | Calcium reversal potential channels | 140mV |
| $a_M$ | Rate constant of sodium channels (closed to open) | 22ms<sup>-1</sup> |
| $b_M$ | Rate constant of sodium channels (open to closed) | 13ms<sup>-1</sup> |
| $a_H$ | Rate constant of sodium channels (non-inactivated to inactivated state) | 0.5ms<sup>-1</sup> |
| $b_H$ | Rate constant of sodium channels (inactivated to non-inactivated state) | 4ms<sup>-1</sup> |
| $a_N$ | Rate constant of sodium channels (closed to open) | 1.5ms<sup>-1</sup> |
| $b_N$ | Rate constant of sodium channels (open to closed) | 0.1ms<sup>-1</sup> |
| $a_Q$ | Rate constant of slow potassium channels (closed to open) | 1.5ms<sup>-1</sup> |
| $b_Q$ | Rate constant of slow potassium channels (open to closed) | 0.025-0.038ms<sup>-1</sup> |
| $a_P$ | Rate constant of calcium channels (closed to open) | 0.008ms<sup>-1</sup> |
| $b_P$ | Rate constant of calcium channels (open to closed) | 0.014 -0.016ms<sup>-1</sup> |

$C_{dendrite} = 2\pi \cdot r_D \cdot l_D \cdot C_m$
$C_{soma} = 2\pi \cdot r_S \cdot l_S \cdot C_m$

**Table S7:**
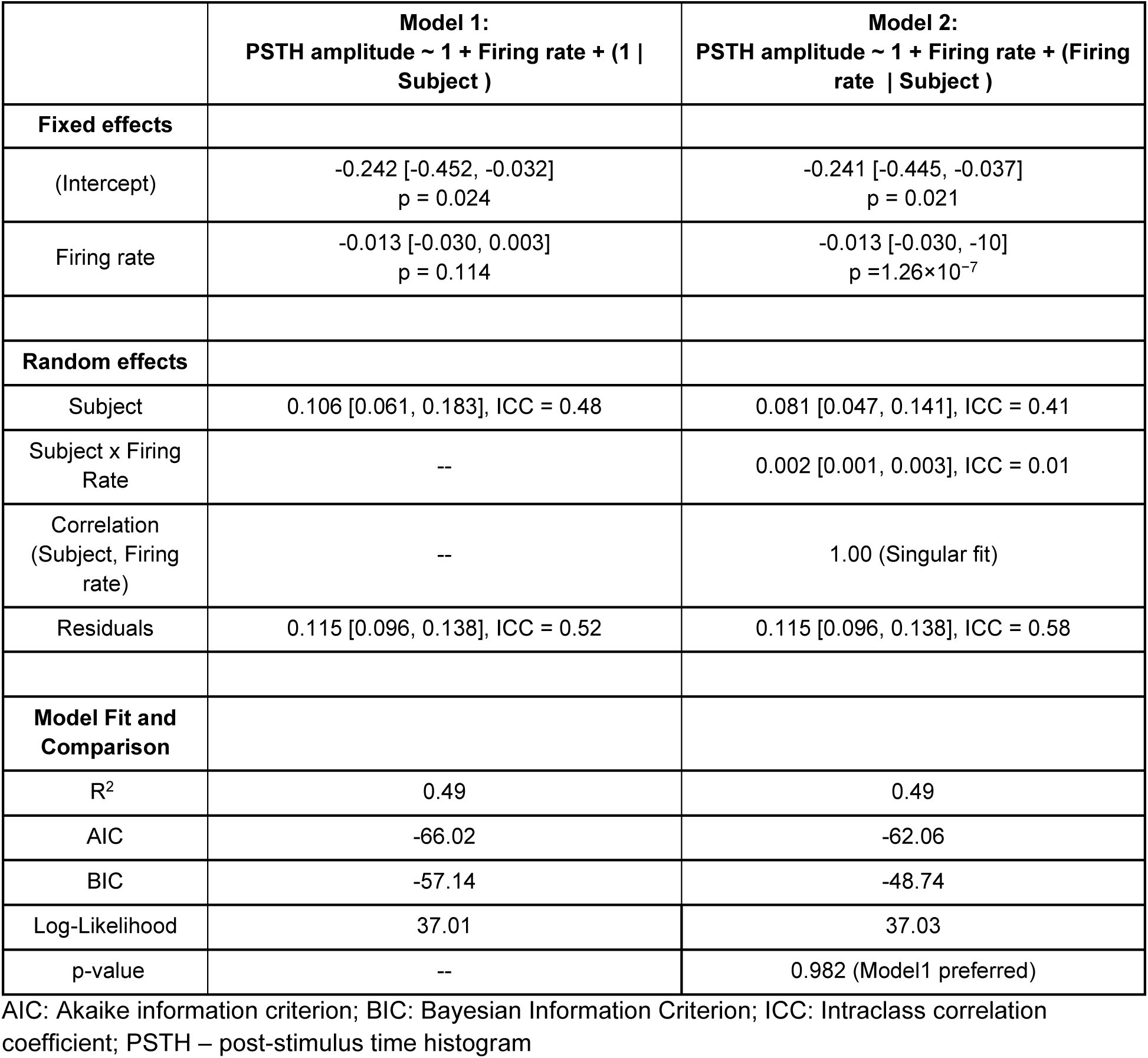
Fixed and random-effects variables and likelihood ratio test from random intercept (model 1) and random intercept and random slope (model 2) linear mixed models for the relationship between PSTH inhibition amplitude and firing rate for cutaneous silent period (CSP) measured from first dorsal interosseous (FDI) muscle

**Table S8:** Fixed and random-effects variables and likelihood ratio test from random intercept (model 1) and random intercept and random slope (model 2) linear mixed models for the relationship between PSF inhibition amplitude and firing rate for cutaneous silent period (CSP) measured from first dorsal interosseous (FDI) muscle

|  | **Model 1:**<br>PSF amplitude ~ 1 + Firing rate + (1 \| Subject ) | **Model 2:**<br>PSF amplitude ~ 1 + Firing rate + (Firing rate \| Subject ) |
| --- | --- | --- |
| **Fixed effects** |  |  |
| (Intercept) | -111 [-203, -21]<br>p = 0.016 | -111 [-254, 31]<br>p = 0.122 |
| Firing rate | -4 [-11, 3]<br>p = 0.234 | -4 [-15, 6]<br>p = 0.384 |
| **Random effects** |  |  |
| Subject | 63 [37, 105], ICC = 0.58 | 0.178 [65, 492], ICC = 0.76 |
| Subject x Firing Rate | -- | 12 [4, 38], ICC = 0.05 |
| Correlation (Subject, Firing rate) | -- | -0.93 [-0.99, -0.45] |
| Residuals | 46 [39, 56], ICC = 0.42 | 43 [35, 53], ICC = 0.18 |
| **Model Fit and Comparison** |  |  |
| R<sup>2</sup> | 0.68 | 0.74 |
| AIC | 755.05 | 758.03 |
| BIC | 763.93 | 771.34 |
| Log-Likelihood | -373.52 | -373.01 |
| p-value | -- | 0.600 (Model1 preferred) |

AIC: Akaike information criterion; BIC: Bayesian Information Criterion; ICC: Intraclass correlation coefficient; PSF – peristimulus frequencygram

**Table S9:**
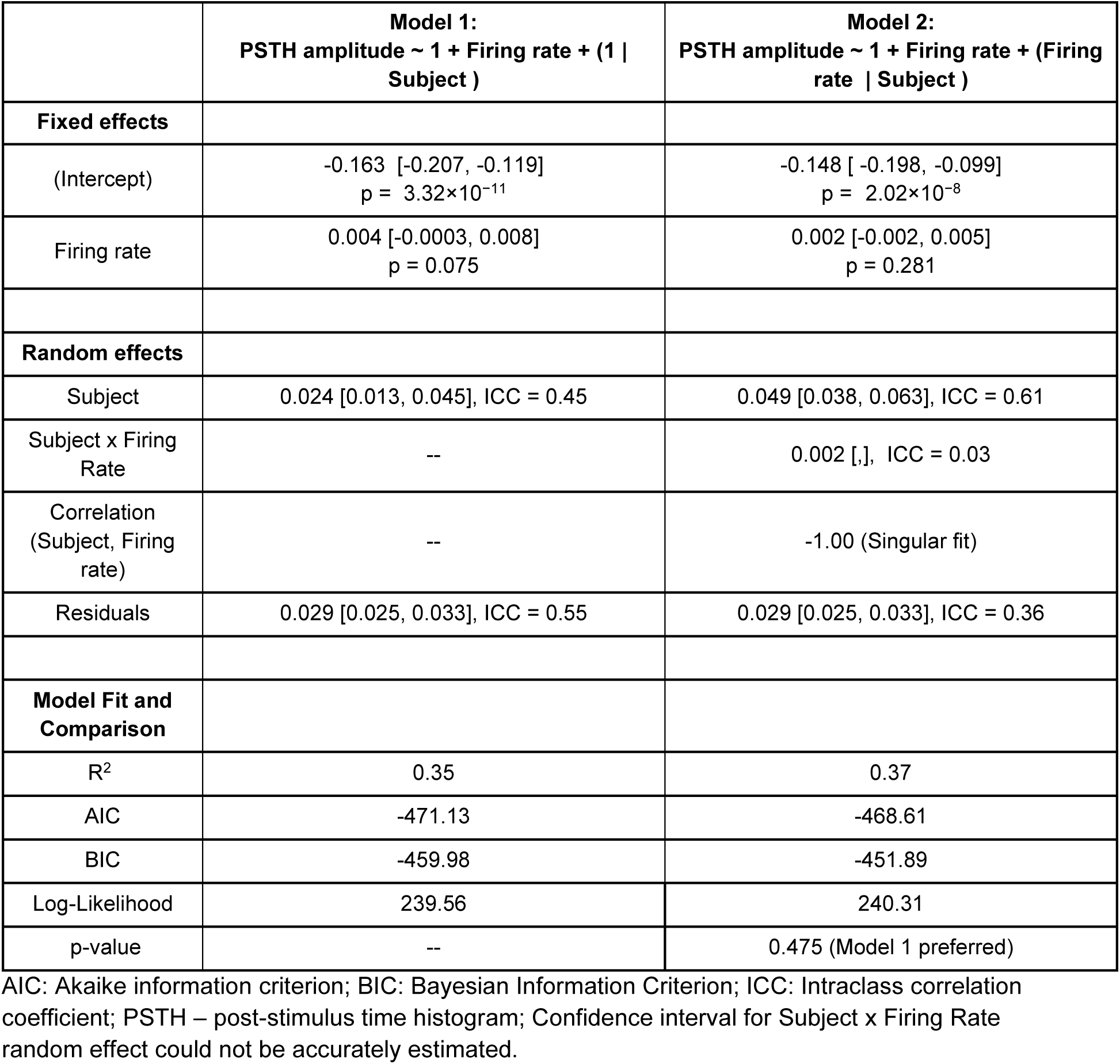
Fixed and random-effects variables and likelihood ratio test from random intercept (model 1) and random intercept and random slope (model 2) linear mixed models for the relationship between PSTH inhibition amplitude and firing rate for reciprocal inhibition measured from tibialis anterior (TA) muscle

**Table S10:** Fixed and random-effects variables and likelihood ratio test from random intercept (model 1) and random intercept and random slope (model 2) linear mixed models for the relationship between PSF inhibition amplitude and firing rate for reciprocal inhibition measured from tibialis anterior (TA) muscle

|  | **Model 1:**<br>PSF amplitude ~ 1 + Firing rate + (1 \| Subject ) | **Model 2:**<br>PSF amplitude ~ 1 + Firing rate + (Firing rate \| Subject ) |
| --- | --- | --- |
| **Fixed effects** |  |  |
| (Intercept) | 17 [ -3 , 39]<br>p = 0.101 | -9 [ -38 , 21]<br>p = 0.570 |
| Firing rate | -4 [-6, -2]<br>p = 1.70×10<sup>-4</sup> | -1 [-5, 3]<br>p = 0.766 |
| **Random effects** |  |  |
| Subject | 15 [9, 27], ICC = 0.54 | 30 [11, 84], ICC = 0.64 |
| Subject x Firing Rate | -- | 5 [2, 10], ICC = 0.11 |
| Correlation (Subject, Firing rate) | -- | -1 [Singular fit] |
| Residuals | 13 [11, 15], ICC = 0.46 | 12 [11, 14], ICC = 0.25 |
| **Model Fit and Comparison** |  |  |
| R<sup>2</sup> | 0.73 | 0.76 |
| AIC | 993.98 | 982.52 |
| BIC | 1005.10 | 999.24 |
| Log-Likelihood | -492.99 | -485.26 |
| p-value | -- | 4.38×10<sup>-4</sup> (Model 2 preferred) |

AIC: Akaike information criterion; BIC: Bayesian Information Criterion; ICC: Intraclass correlation coefficient; PSF – peristimulus frequencygram

**Table S11:** Fixed and random-effects variables and likelihood ratio test from random intercept (model 1) and random intercept and random slope (model 2) linear mixed models for the simulated data pertaining to the LM_sim_ chosen for each subject, showing the relationship between Inhibition duration and firing rate for simulated cutaneous silent period (CSP)

|  | **Model 1:**<br>**Inhibition Duration ~ 1 + Firing rate + (1 \| Subject )** | **Model 2:**<br>**Inhibition Duration ~ 1 + Firing rate + (Firing rate \| Subject )** |
| --- | --- | --- |
| **Fixed effects** |  |  |
| (Intercept) | 288 ms [265, 311]<br>p = $1.97 \times 10^{-50}$ | 289 ms [251, 328]<br>p = $7.65 \times 10^{-30}$ |
| Firing rate | -14 ms [-16, -12]<br>p = $8.73 \times 10^{-31}$ | -14 ms [-27, -11]<br>p = $2.85 \times 10^{-17}$ |
| **Random effects** |  |  |
| Subject | 17 [10, 28], ICC = 0.57 | 53 [28, 98], ICC = 0.77 |
| Subject x Firing Rate | -- | 4 [2, 7], ICC = 0.06 |
| Correlation (Subject, Firing rate) | -- | -0.96 [-0.99, -0.80] |
| Residuals | 13 [11, 15], ICC = 0.43 | 12 [11, 14], ICC = 0.17 |
| **Model Fit and Comparison** |  |  |
| R<sup>2</sup> | 0.77 | 0.80 |
| AIC | 1090.9 | 1086.9 |
| BIC | 1102.4 | 1104.2 |
| Log-Likelihood | -541.45 | -537.46 |
| p-value | -- | 0.019 (Model 2 preferred) |

AIC: Akaike information criterion; BIC: Bayesian Information Criterion; ICC: Intraclass correlation coefficient

**Table S12:** Fixed and random-effects variables and likelihood ratio test from random intercept (model 1) and random intercept and random slope (model 2) linear mixed models for the simulated data pertaining to the LM_sim_ chosen for each subject, showing the relationship between PSTH inhibition amplitude and firing rate for simulated cutaneous silent period (CSP)

|  | **Model 1:**<br>**PSTH amplitude ~ 1 + Firing rate + (1 \|**<br>**Subject )** | **Model 2:**<br>**PSTH amplitude ~ 1 + Firing rate + (Firing**<br>**rate \| Subject )** |
| --- | --- | --- |
| **Fixed effects** |  |  |
| (Intercept) | -0.532 [ -0.655, -0.408]<br>p = 3.84×10<sup>-14</sup> | -0.538 [-0.685, -0.390]<br>p = 3.84×10<sup>-11</sup> |
| Firing rate | 0.007 [0.002, 0.012]<br>p = 0.009 | 0.008 [0.0004, 0.015]<br>p = 0.036 |
| **Random effects** |  |  |
| Subject | 0.168 [0.106, 0.268], ICC = 0.80 | 0.212 [0.120, 0.373], ICC = 0.95 |
| Subject x Firing<br>Rate | -- | 0.002 [0.001, 0.003], ICC = 0.01 |
| Correlation<br>(Subject, Firing<br>rate) | -- | -0.64 [-0.94, 0.22] |
| Residuals | 0.041 [0.035, 0.045], ICC = 0.20 | 0.009 [0.002, 0.021], ICC = 0.03 |
| **Model Fit and**<br>**Comparison** |  |  |
| R<sup>2</sup> | 0.94 | 0.95 |
| AIC | -400.82 | -399.26 |
| BIC | -389.32 | -382.01 |
| Log-Likelihood | 204.41 | 205.63 |
| p-value | -- | 0.296 (Model1 preferred) |

AIC: Akaike information criterion; BIC: Bayesian Information Criterion; ICC: Intraclass correlation coefficient; PSTH – post-stimulus time histogram

**Table S13:** Fixed and random-effects variables and likelihood ratio test from random intercept (model 1) and random intercept and random slope (model 2) linear mixed models for the simulated data pertaining to the LM_sim_ chosen for each subject, showing the relationship between PSF inhibition amplitude and firing rate for simulated cutaneous silent period (CSP)

|  | **Model 1:**<br>**PSF amplitude ~ 1 + Firing rate + (1 \| Subject )** | **Model 2:**<br>**PSF amplitude ~ 1 + Firing rate + (Firing rate \| Subject )** |
| --- | --- | --- |
| **Fixed effects** |  |  |
| (Intercept) | -98 [-150, -48]<br>p = $1.94 \times 10^{-4}$ | -99 [-178, -19]<br>p = 0.015 |
| Firing rate | -6 [-9, -3]<br>p = $4.24 \times 10^{-4}$ | -6 [-12, 0.3]<br>p = 0.064 |
| **Random effects** |  |  |
| Subject | 54 [34, 87], ICC = 0.69 | 115 [65, 204], ICC = 0.79 |
| Subject x Firing Rate | -- | 9 [5, 16], ICC = 0.06 |
| Correlation (Subject, Firing rate) | -- | -0.88 [-0.97, -0.55] |
| Residuals | 24 [21, 27], ICC = 0.31 | 21 [19, 24], ICC = 0.14 |
| **Model Fit and Comparison** |  |  |
| R<sup>2</sup> | 0.84 | 0.87 |
| AIC | 1257.4 | 1247.7 |
| BIC | 1268.9 | 1265.0 |
| Log-Likelihood | -624.72 | -617.87 |
| p-value | -- | 0.001 (Model 2 preferred) |

AIC: Akaike information criterion; BIC: Bayesian Information Criterion; ICC: Intraclass correlation coefficient; PSF – peristimulus frequencygram

**Table S14:**
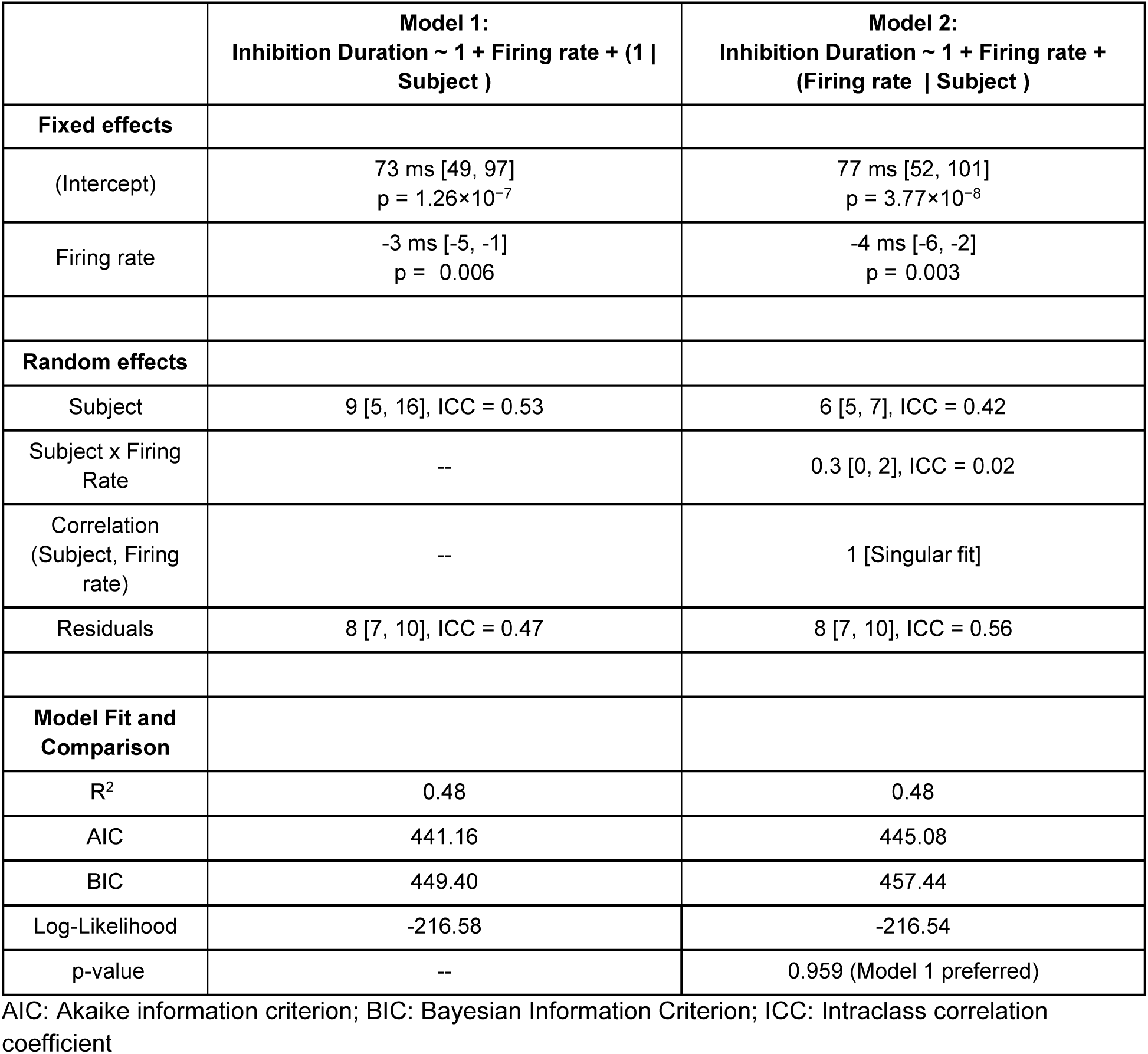
Fixed and random-effects variables and likelihood ratio test from random intercept (model 1) and random intercept and random slope (model 2) linear mixed models for the simulated data pertaining to the LM_sim_ chosen for each subject, showing the relationship between Inhibition Duration and firing rate for simulated reciprocal inhibition

|  | **Model 1:**<br>**Inhibition Duration ~ 1 + Firing rate + (1 \|**<br>**Subject )** | **Model 2:**<br>**Inhibition Duration ~ 1 + Firing rate +**<br>**(Firing rate \| Subject )** |
| --- | --- | --- |
| **Fixed effects** |  |  |
| (Intercept) | 73 ms [49, 97]<br>p = 1.26×10<sup>-7</sup> | 77 ms [52, 101]<br>p = 3.77×10<sup>-8</sup> |
| Firing rate | -3 ms [-5, -1]<br>p = 0.006 | -4 ms [-6, -2]<br>p = 0.003 |
| **Random effects** |  |  |
| Subject | 9 [5, 16], ICC = 0.53 | 6 [5, 7], ICC = 0.42 |
| Subject x Firing<br>Rate | -- | 0.3 [0, 2], ICC = 0.02 |
| Correlation<br>(Subject, Firing<br>rate) | -- | 1 [Singular fit] |
| Residuals | 8 [7, 10], ICC = 0.47 | 8 [7, 10], ICC = 0.56 |
| **Model Fit and**<br>**Comparison** |  |  |
| R<sup>2</sup> | 0.48 | 0.48 |
| AIC | 441.16 | 445.08 |
| BIC | 449.40 | 457.44 |
| Log-Likelihood | -216.58 | -216.54 |
| p-value | -- | 0.959 (Model 1 preferred) |

AIC: Akaike information criterion; BIC: Bayesian Information Criterion; ICC: Intraclass correlation coefficient

**Table S15:** Fixed and random-effects variables and likelihood ratio test from random intercept (model 1) and random intercept and random slope (model 2) linear mixed models for the simulated data pertaining to the LM_sim_ chosen for each subject, showing the relationship between PSTH inhibition amplitude and firing rate for simulated reciprocal inhibition

|  | **Model 1:**<br>**PSTH amplitude ~ 1 + Firing rate + (1 \|**<br>**Subject )** | **Model 2:**<br>**PSTH amplitude ~ 1 + Firing rate + (Firing**<br>**rate \| Subject )** |
| --- | --- | --- |
| **Fixed effects** |  |  |
| (Intercept) | -0.085 [-0.153, -0.017]<br>p = 0.014 | -0.096 [-0.174, -0.018]<br>p = 0.015 |
| Firing rate | -0.002 [-0.009, 0.003]<br>p = 0.358 | -0.002 [-0.010, 0.006]<br>p = 0.653 |
| **Random effects** |  |  |
| Subject | 0.028 [0.016, 0.051], ICC = 0.56 | 0.041 [.,], ICC = 0.59 |
| Subject x Firing<br>Rate | -- | 0.007 [0.005, 0.008], ICC = 0.10 |
| Correlation<br>(Subject, Firing<br>rate) | -- | -1.00 (Singular fit) |
| Residuals | 0.022 [ 0.018, 0.027], ICC = 0.44 | 0.022 [0.018, 0.026], ICC = 0.31 |
| **Model Fit and**<br>**Comparison** |  |  |
| R<sup>2</sup> | 0.60 | 0.62 |
| AIC | -240.93 | -239.01 |
| BIC | -232.69 | -226.65 |
| Log-Likelihood | 124.47 | 125.51 |
| p-value | -- | 0.352 (Model 1 preferred) |

AIC: Akaike information criterion; BIC: Bayesian Information Criterion; ICC: Intraclass correlation coefficient; PSTH – post-stimulus time histogram; Confidence interval for Subject in Model 2 could not be accurately estimated.

**Table S16:** Fixed and random-effects variables and likelihood ratio test from random intercept (model 1) and random intercept and random slope (model 2) linear mixed models for the simulated data pertaining to the LM_sim_ chosen for each subject, showing the relationship between PSF inhibition amplitude and firing rate for simulated reciprocal inhibition

|  | **Model 1:**<br>PSF amplitude ~ 1 + Firing rate + (1 \| Subject ) | **Model 2:**<br>PSF amplitude ~ 1 + Firing rate + (Firing rate \| Subject ) |
| --- | --- | --- |
| **Fixed effects** |  |  |
| (Intercept) | -4 [-35, 27]<br>p = 0.786 | -1 [-32, 29]<br>p = 0.928 |
| Firing rate | -2 [-5, 1]<br>p = 0.254 | -2 [-5, 1]<br>p = 0.169 |
| **Random effects** |  |  |
| Subject | 6 [2, 14], ICC = 0.33 | 11 [8, 15], ICC = 0.47 |
| Subject x Firing Rate | -- | 0.5 [0.3, 0.7], ICC = 0.02 |
| Correlation (Subject, Firing rate) | -- | -1 [Singular fit] |
| Residuals | 12 [10, 14], ICC = 0.67 | 12 [10, 14], ICC = 0.51 |
| **Model Fit and Comparison** |  |  |
| R<sup>2</sup> | 0.18 | 0.18 |
| AIC | 471.94 | 475.87 |
| BIC | 480.18 | 488.23 |
| Log-Likelihood | -231.97 | -231.94 |
| p-value | -- | p= 0.966(Model 1 preferred) |

AIC: Akaike information criterion; BIC: Bayesian Information Criterion; ICC: Intraclass correlation coefficient; PSF – peristimulus frequencygram

**Figure S1.**
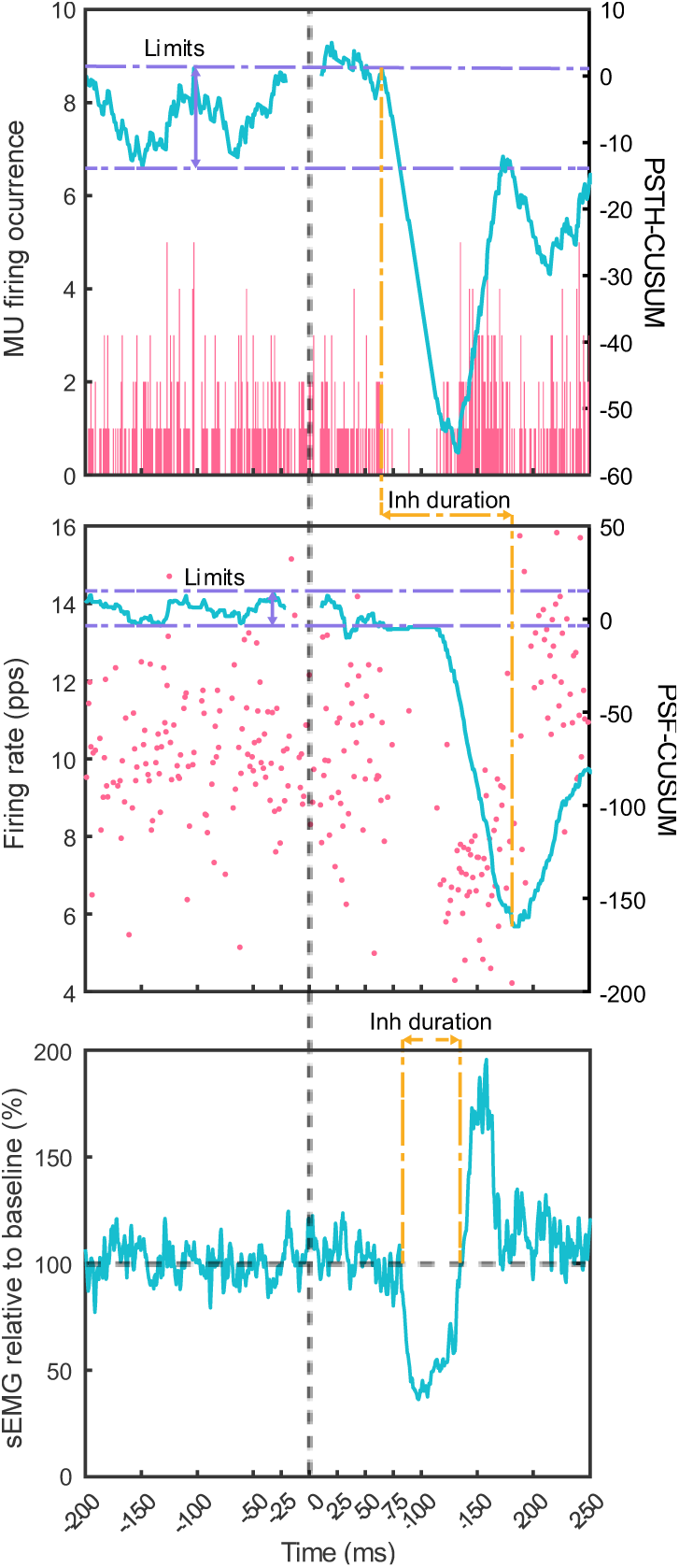
Motor unit firings: PSTH and PSF with CUSUM analysis. Example of CSP motor unit firings with PSTH (top) and PSF (middle) and respective CUSUM, showing the defined start and end of inhibition (dotted yellow lines) and CUSUM limits (dotted purple line). Bottom trace displays the sEMG obtained from the same recording and the estimate of inhibition duration. Black vertical dotted line denotes onset of nerve stimulation.

**Figure S2.**
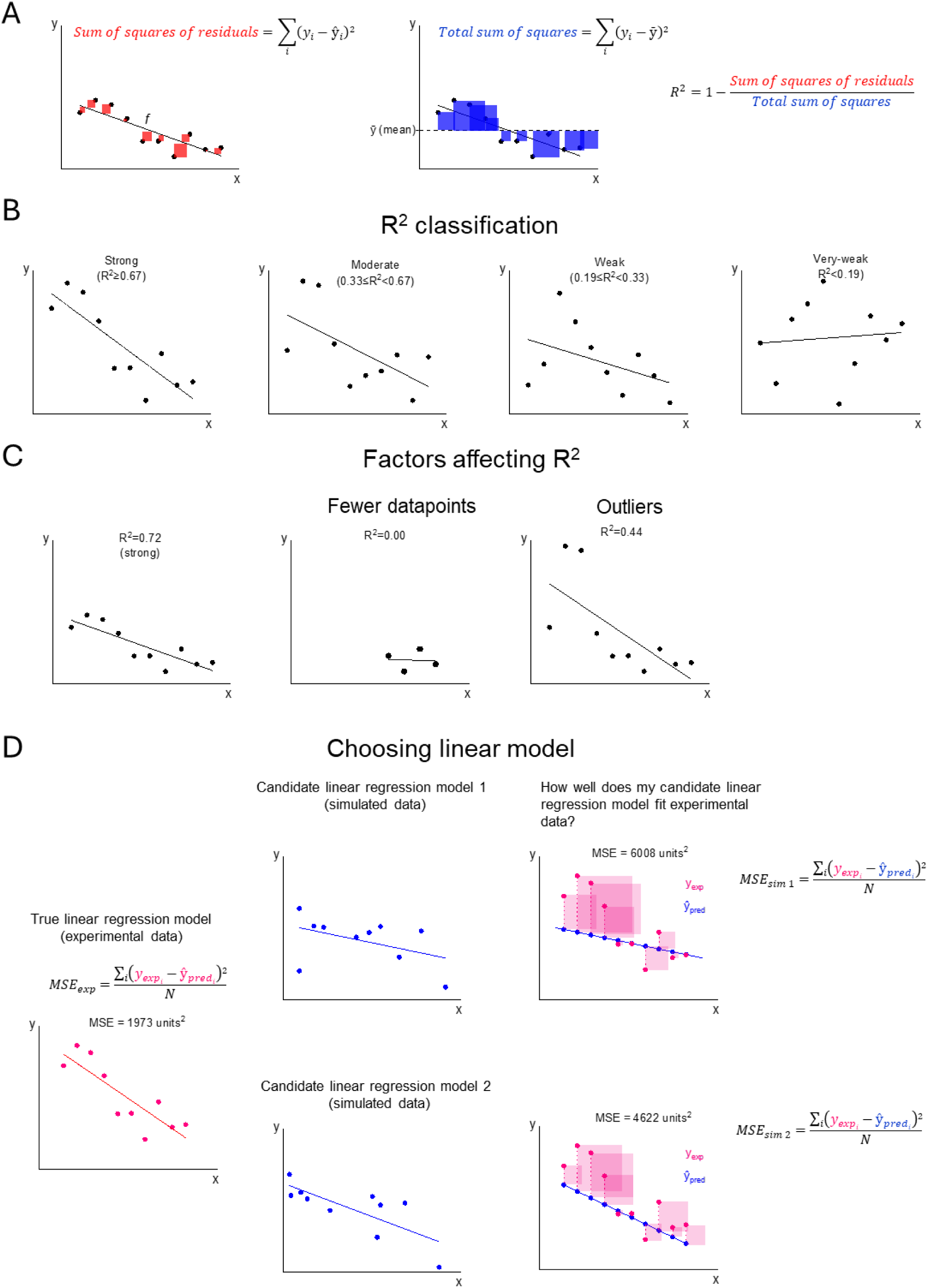
Statistical assessment of variability and model selection in linear regression. **(A)** example of datapoints and respective linear fit (*f*), illustrating the sum of squares of residuals (red, left) and total sum of squares (blue, middle) used to estimate the coefficient of determination (R^2^; right). **(B)** examples of R^2^ of different strengths: strong, moderate, weak and very weak R^2^; **(C)** Illustration of factors influencing the R^2^ such as sample size and datapoint outliers; **(D)** Steps for selecting the best linear model from simulated data: 1) experimental data is used to establish a reference linear model (pink, left), where the mean squared error (MSE) quantifies the true relationship between the predictor (x) and the dependent variable (y); 2) simulated datasets generate multiple candidate regression models (blue), each with its own independent and dependent variables; 3) the linear fit from each simulated model is evaluated against the experimentally measured predictor, with the best linear model selected by prioritizing the one that minimizes the MSE.

**Figure S3.**
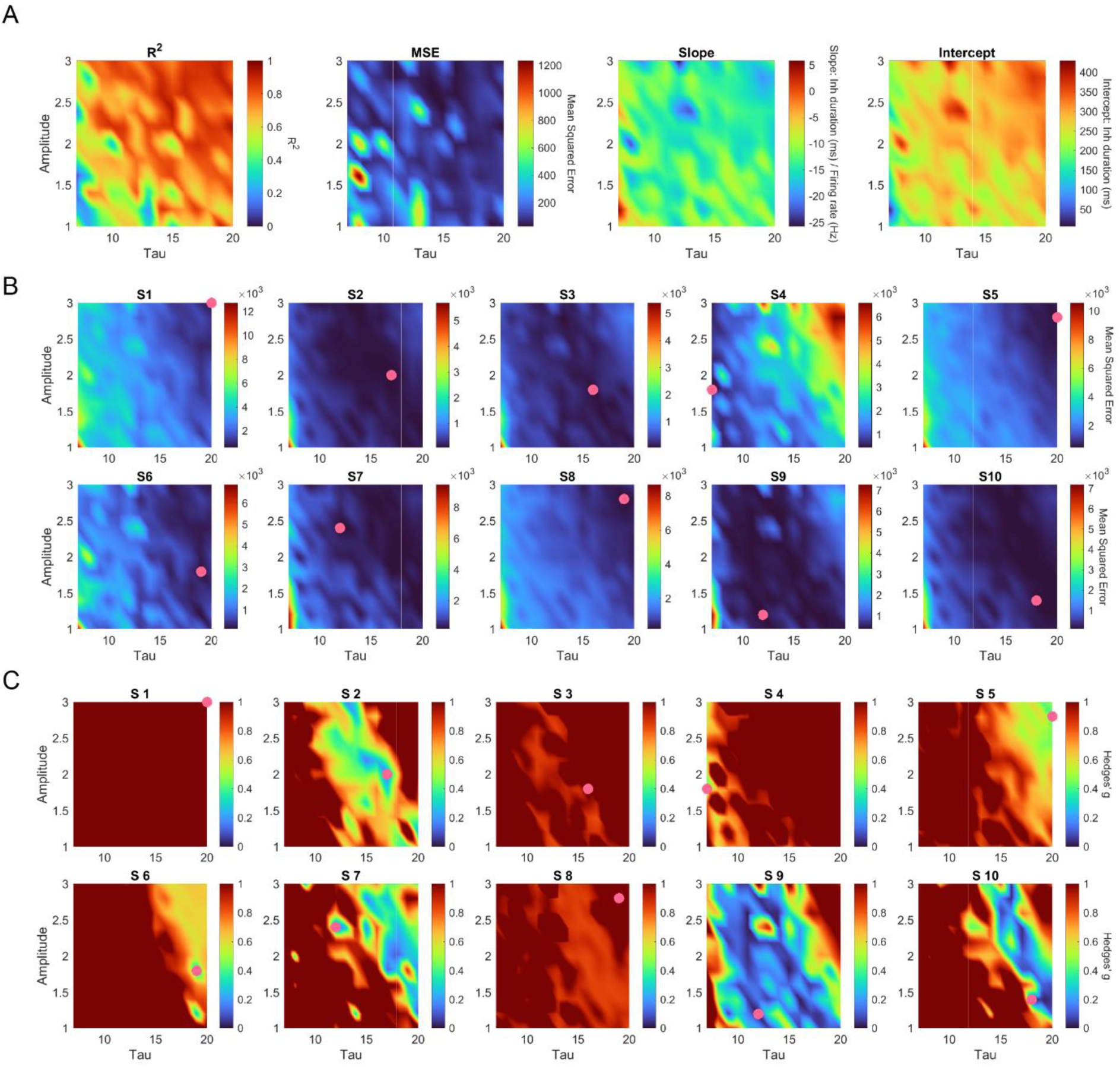
Heatmap and optimization analysis of R^2^ for CSP linear models from an *in silico* biophysical model. **(A)** Heatmaps depicting the R^2^ (left), Mean Squared Error (MSE, middle left), slope (middle right) and intercept (right) of linear regressions for all combinations of amplitude (1–3 a.u.) and duration (τ, 7–20ms) across 154 realizations, with colour-coded intensity (blue to red). **(B, C)** Hyperparameter optimization plots for each of the 10 subjects, illustrating **(B)** Mean Squared Error values and **(C)** Hedges’ *g* for the square of residuals comparisons between LM_exp_ and linear fits obtains obtained across amplitude and τ variations, highlighting the best-fit LM_sim_ selected for validation against CSP HDsEMG data (pink dot).

**Figure S4.**
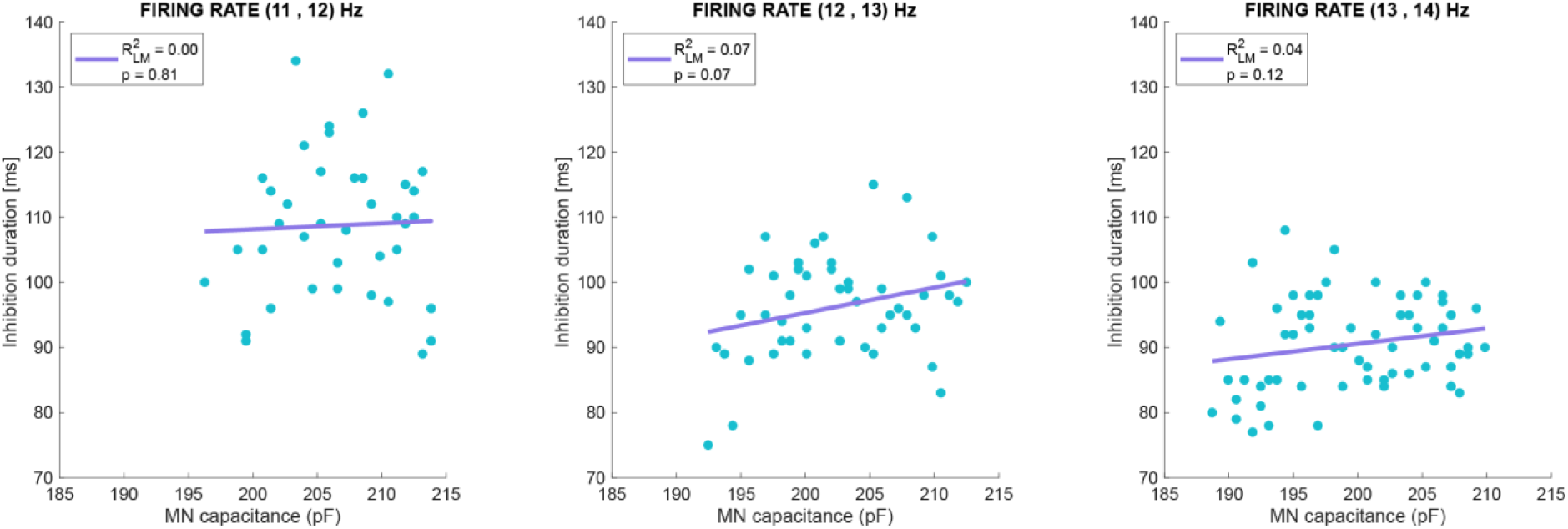
Influence of motoneuron size on inhibition duration *in silico* (optimized parameters for subject 4). Inhibition duration across simulated motoneurons with varying size (blue dots) with each plot representing a different firing rate, with purple line representing the linear regression between capacitance and inhibition duration. Parameters include an inhibitory input amplitude of 1.8a.u. and duration of 7ms.

**Figure S5.**
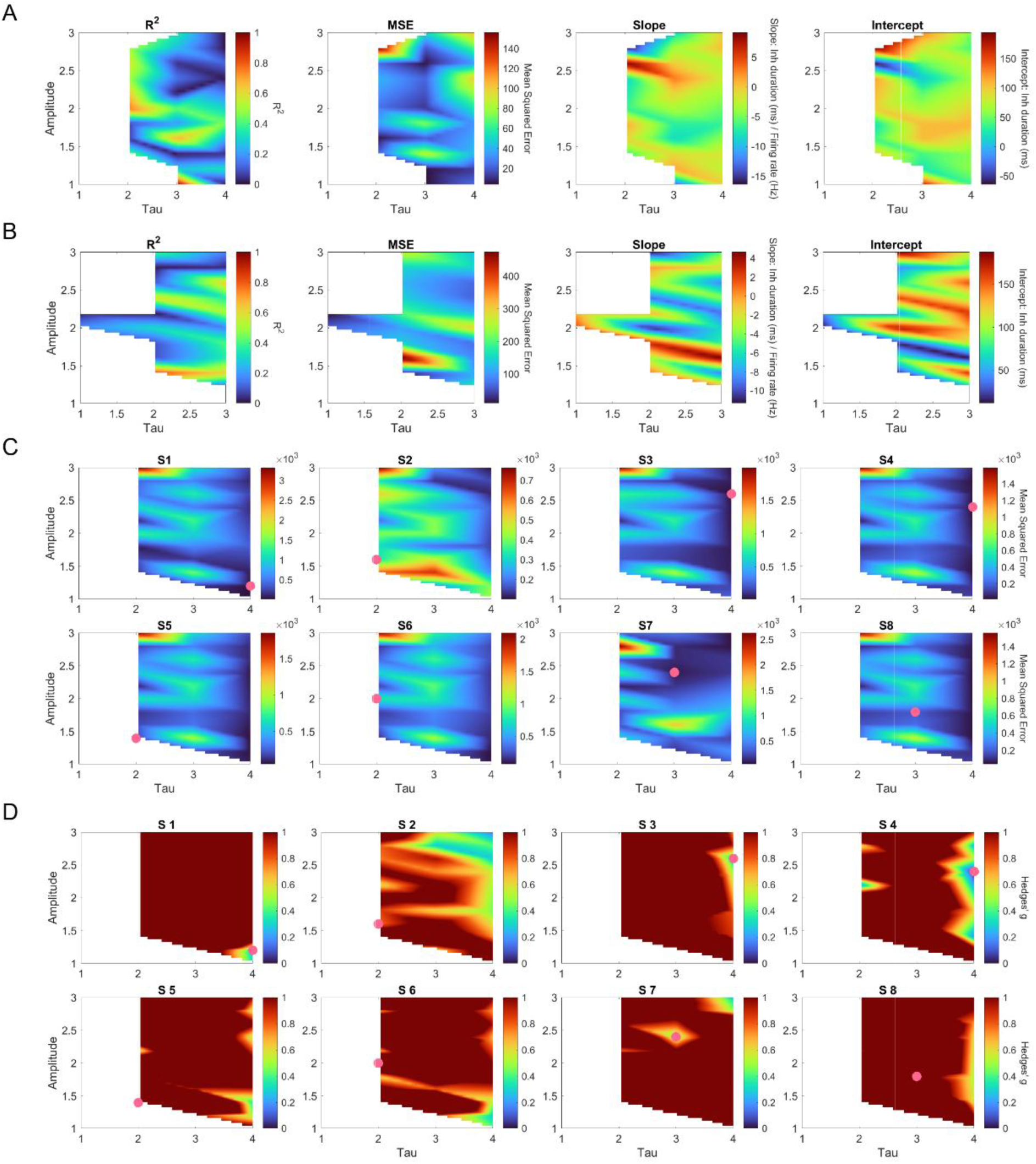
Heatmap and optimization analysis of R^2^ for reciprocal inhibition linear fits obtained through *in silico* biophysical modelling. **(A-B)** Heatmaps depicting the R^2^ (left), Mean Squared Error (MSE, middle left), slope (middle right) and intercept (right) of linear regressions for all combinations of amplitude (1–3a.u.) and duration (τ, 1–4ms) across 39 realizations with excitatory common input of **(A)** 0.15a.u. and across 33 realizations with **(B)** excitatory common 0.2a.u., with colour-coded intensity (blue to red). **(C, D)** Hyperparameter optimization plots for each of the 8 subjects, illustrating **(C)** Mean Squared Error values and **(D)** Hedges’ *g* for the square of residuals comparisons between LM_exp_ and linear fits obtains obtained across amplitude and τ variations, highlighting the best-fit LM_sim_ selected for validation against reciprocal inhibition HDsEMG data (pink dot). All *in silico* models except for S2 and S7 received 0.15a.u. of excitatory common input.

